# KODAMA enables self-guided weakly supervised learning in spatial transcriptomics

**DOI:** 10.1101/2025.05.28.656544

**Authors:** Ebtesam A. Abdel-Shafy, Moussa Kassim, Alessia Vignoli, Farag Mamdouh, Svitlana Tyekucheva, Dalia Ahmed, Dupe Ojo, Brendon Price, Fabio Socciarelli, Nancy Paola Duarte-Delgado, Martin Ocharo, Chiamaka Jessica Okeke, Luca Triboli, David A. MacIntyre, Massimo Loda, Devanand Sarkar, Dinesh Gupta, Silvano Piazza, Luiz Fernando Zerbini, Leonardo Tenori, Stefano Cacciatore

## Abstract

Spatial transcriptomics provides researchers with a powerful tool to investigate gene expression patterns within their native positional context in tissues, revealing intricate cellular relationships. However, analyzing spatial transcriptomics data presents unique challenges due to their high dimensionality and complexity. Several competitive tools have emerged, each aiming to integrate spatial dependencies into their workflow. Yet, the overall performance of these approaches remains constrained by limitations in prediction accuracy and compatibility with specific data types. Here, we introduce the third version of the KODAMA algorithm, specifically tailored for spatial transcriptomics analysis. This upgraded version incorporates a novel approach that effectively reduces data dimensionality while preserving spatial information. At its core, the KODAMA algorithm employs parallel iterations that enforce spatial constraints throughout an embedded clustering process. Our method provides the flexibility to simultaneously analyze multiple samples, streamlining analysis workflows. Additionally, it extends beyond traditional 2D analysis to accommodate multidimensional datasets, including 3D spatial information. KODAMA seamlessly integrates into various pipelines, such as Seurat and Giotto. Extensive evaluations of KODAMA on 10x Visium, Visium HD, and image-based 3D datasets demonstrate its high accuracy in spatial domain prediction. Comparative analyses against alternative dimensionality reduction techniques and spatial analysis tools consistently validate and highlight KODAMA’s superior performance in unraveling the spatial organization of cellular components across both single and integrated tissue samples.

## Main

Multicellular organisms, including humans, boast a remarkable diversity of tissues and organs, each comprised of highly complex specialized cell types. Despite sharing a common genetic blueprint, each cell type exhibits unique gene expression profiles shaped by intrinsic regulatory mechanisms and extrinsic signals from the surrounding microenvironment [1, 2]. High-throughput transcriptomic technologies, particularly single-cell RNA sequencing (scRNA-seq), have illuminated previously obscured cellular heterogeneity and regulatory networks. However, scRNA-seq is limited by its inability to preserve spatial context, as tissue dissociation during preparation disrupts the native cellular architecture [3, 4]. This limitation poses challenges, especially for complex tissues like the nervous system, where spatial organization is critical for proper function [5, 6], and tumors, characterized by a heterogenous microenvironment with extensively variable cellular niches [7]. In response to these challenges, spatial transcriptomics (ST) has emerged as a powerful solution, allowing researchers to preserve spatial information while profiling gene expression within intact tissue samples with a deeper understanding of cellular interactions and tissue structure [8–11]. Each ST approach offers distinct advantages and trade-offs in spatial resolution, throughput, sensitivity, and scalability, catering to different experimental needs and biological contexts [12].

Image-based ST (*i*ST) techniques expand and scale the principles of fluorescence *in situ* hybridization (FISH) to simultaneously detect multiple transcripts while preserving spatial resolution. These techniques include multiplexed error-robust FISH (MERFISH) [13], sequential FISH (seqFISH) [14], and spatially-resolved transcript amplicon readout mapping (STARmap) [15]. Despite the relatively low coverage of *i*ST, capturing only a few hundred to a few thousand distinct genes, these techniques achieve high transcript depth, indicating a substantial number of unique RNA molecules captured for each gene. Integrating spatial information across multiple tissue slices enables the reconstruction of three-dimensional (3D) gene expression maps, revealing tissue architecture and cellular interactions with high clarity [16, 17]. Advanced techniques like 3D MERFISH enhance the exploration of gene expression patterns across tissue depth, with enhanced resolution and sensitivity [18].

In contrast, array-based ST (*a*ST) platforms, including Visium [19], Slide-Seq [20, 21], and Stereo-Seq [22], operate on a mini-bulk transcriptomics principle, where data are aggregated at defined spatial units (spots), which represent areas of the tissue at multicellular level. Techniques like Visium employ slides embedded with arrayed oligo-dT spots to capture and spatially barcode poly(A) RNAs, profiled using next-generation sequencing. This process can be integrated with hematoxylin and eosin (H&E) staining or limited antigen-antibody staining on the same fresh-frozen or formalin-fixed paraffin-embedded (FFPE) tissue section. While they offer valuable insights into spatial gene expression, their resolution may limit the ability to discern subtle spatial variations within tissues. Nonetheless, advancements like 10x VisiumHD are designed to allow users to comprehensively characterize FFPE tissue sections by measuring whole transcriptome spatial gene expression at single-cell resolution. This new platform can profile 11 million spots in a continuous grid pattern of 2 μm squares, compared to just 5,000 spots detectable in a hexagonal arrangement of 55 μm spots of its predecessor.

Over time, spatial transcriptomics technologies have expanded, generating substantial amounts of unstructured data containing spatial and gene expression information [23]. This growth has prompted the development of computational methods for analyzing data from multiple samples. Capturing 3D spatial information introduces additional complexity, necessitating the development of specialized tools for effective interpretation. Furthermore, the increased resolution of spatial transcriptomics (ST) techniques, particularly platforms like VisiumHD, introduces challenges in data processing and computational efficiency. As data scale grows, novel approaches become essential for managing and interpreting this vast amount of information.

Most existing methods for detecting spatial regions are based on partitional clustering algorithms, such as BayesSpace [24]and BASS[25]. Partitional clustering reduces the information into non-overlapping groups or clusters, where each data point is assigned to one cluster. As such, these methods do not capture the relative distances or relationships between groups or account for potential subclassifications or hierarchical structures within the data, potentially losing valuable insights about the underlying patterns and connections. Dimensionality reduction methods help to reveal underlying features [26]. While algorithms such as *t*-stochastic neighbor embedding (*t*-SNE) [27] and Uniform Manifold Approximation and Projection (UMAP) [28] are widely used for single-cell transcriptomic data, they lack specific capabilities for handling tissue spatial coordinates. Recently, PRECAST [29] and BANKSY [30] have been proposed for the analysis of ST datasets by integrating spatial information into a preprocessing step followed by a dimensionality reduction workflow utilizing *t-*SNE and UMAP, respectively. Despite their strengths, both methods have limitations in generalizability, as their performance may vary significantly across different ST datasets. The limitations of current methods underscore the need for further development of more robust and comprehensive computational methods.

Here, we present the third version of the KODAMA algorithm—now spatially informed—to explore the complex interplay between gene expression patterns and spatial organization within tissues. KODAMA [31] is a hybrid algorithm that transitions from unsupervised learning to self-supervised refinement through iterative cross-validation, ultimately incorporating weak supervision via spatial constraints. Given its progressive learning nature, we can define this as a new class of machine learning algorithms, self-guided weakly supervised learning. By integrating spatial coordinates into the analysis, KODAMA provides new insights into tissue architecture using datasets generated on different platforms. Benchmarking demonstrates that it outperforms other dimensionality reduction methods and ST dataset clustering tools, particularly in 3D and multi-sample contexts. It also revealed new insights in the large-scale VisiumHD dataset, where data size and computational complexity are significant obstacles. Moreover, its compatibility and seamless integration with widely used pipelines — including Seurat [32], Giotto [33], and SpatialExperiment [34] — ensures effortless integration into existing workflows, fostering high-resolution, spatially informed biological discoveries.

## Results

### ST data analysis using KODAMA

KODAMA is a machine-learning algorithm for feature extraction from noisy and high-dimensional data [31, 35]. Similar to self-supervised learning methods, where clustering can be used to generate supervised models, we previously introduced the concept that clustering itself can be improved by editing the class labels of samples not correctly predicted in cross-validation [31]. This builds from the previous work, showing that class prediction can be refined by removing samples incorrectly predicted in cross-validation [36]. In KODAMA, class labels are iteratively refined through a cross-validation procedure in which misclassified labels are replaced with their predicted class values. Depending on the reassignment, a higher cross-validated accuracy is achieved (**Fig. 1a**). This procedure closely aligns with the refinement process in weakly supervised learning algorithms, where iterative corrections enhance classification accuracy despite incomplete or uncertain labels. Spatial information can be integrated to guide the cross-validation accuracy maximization. At the beginning of each iteration, observations (*e.g.*, samples, spots, or cells) are grouped based on their local proximity. All samples within the same proximity group are forced to share the same class label during the iterative process (**Fig. 1b**). This process ensures that, rather than switching individual sample labels, entire proximity groups are reassigned class labels, preserving local spatial relationships. By incorporating spatial information, KODAMA can better capture gene expression patterns within the tissue architecture. To reduce drastically both the space and time complexity associated with analyzing such large data sets, we implemented a novel landmarking procedure where the cross-validation accuracy maximization procedure is performed only on a selected number of samples (landmarks) and the class labels of the remaining samples (projections) are predicted through a supervised model (**Fig. 1c**).

**Fig. 1:**
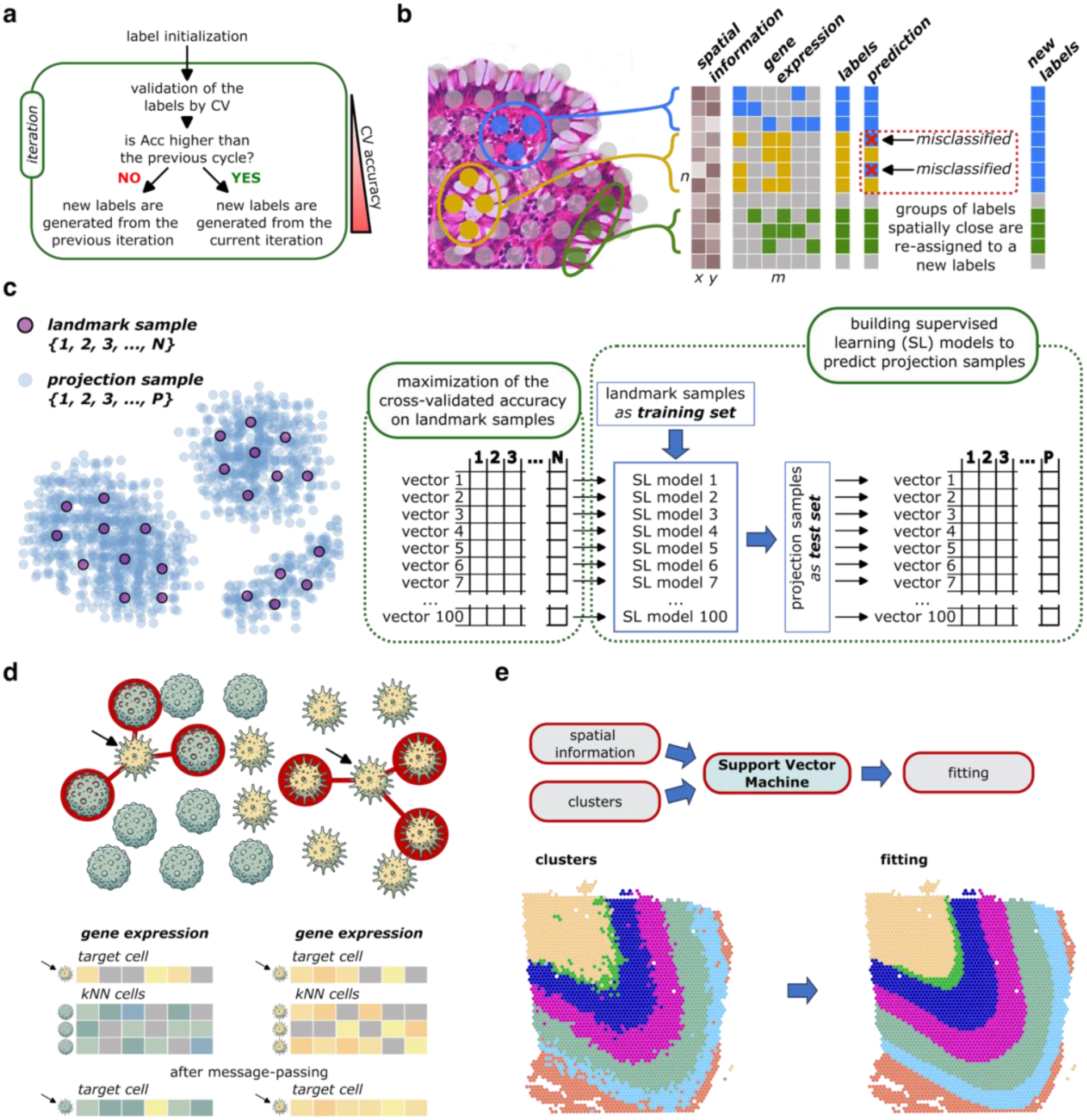
The KODAMA framework. **a,** Simplified diagram showing the iterative procedure that leads to the maximization of the cross-validated accuracy. **b,** Graphical representation of how spatial information is integrated in the maximization of the cross-validated accuracy. Starting from a matrix of m features and n observations, the latter are clustered based on their spatial information (x, y) as the initial step of KODAMA. After running a cross-validation using the labels generated by the clustering, we will obtain a prediction class for each observation. In the spatially informed KODAMA, the misclassified labels are identified, and they drive the change to the class labels of the whole cluster group. **c,** In the landmarking procedure, a limited number (N) of landmarks are chosen among all observations. Independent iterative procedures of cross-validated accuracy maximization are performed using only landmark observations. Resulting vectors generated by the cross-validated accuracy maximization are used to build a supervised model to predict the remaining (P) observations (projections). The landmark vectors and the projection vectors are then combined to build a dissimilarity matrix. The dissimilarity matrix can then be transformed into a dimensional space using UMAP or *t*-SNE. **d,** Message-passing preprocessing combines the profiles of neighboring cells into spatially aggregated expression profiles. The target cell profile is changed to be more similar to the neighboring cells. **e,** The SVM model is built on the observations’ spatial information and the cluster obtained from the KODAMA dimensionality space. SVM refines the results obtained, resulting in well spatially defined clustering.

We have expanded KODAMA into a versatile framework by incorporating additional tools, enhancing its ability to analyze diverse types of ST data with greater depth and flexibility. To improve the detection of tissue microanatomical structures in *i*ST, we developed a simplified preprocessing procedure based on the concept of “message-passing” [37]. By passing information between neighboring spots, the method helps capture the spatial continuity of gene expression, which is crucial for understanding how different regions of the tissue influence each other biologically. This method combines phenotypic information across cells in spatial proximity, with each cell acquiring a portion of its neighbors’ phenotypic information, weighted inversely by distance. The resulting spatially aggregated features reflect both phenotypes and the physical proximity of the cells (**Fig. 1d**).

Gene-gene correlation analysis is crucial for understanding pathway roles within a tissue. However, correlation coefficients, such as Pearson and Spearman, struggle to capture relationships between features in sparse datasets, such as ST data. To address this limitation, correlation tests can be conducted after message-passing (AMP) procedure, which enhances data completeness by leveraging spatial locality to retrieve missing information. This approach ensures a more accurate representation of gene-gene interactions within the tissue architecture.

Tissue microarchitectures can be identified by clustering methods applied to the resulting KODAMA dimensions. To further refine spatial cluster representation, a support vector machine (SVM) model [38] is trained using spatial information and clustering results. The SVM finds optimal separation functions to define clear boundaries between clusters. These refined boundaries enable a clearer visualization of spatial structures (**Fig. 1e**). The spatial resolution of the tissue segmentation can be improved by applying the SVM to a spatial coordinate with higher resolution. Selected tissue segments can be converted into GeoJSON files, making them compatible with microscopy image analysis tools like QuPath [39], facilitating seamless integration with histopathological images.

We applied KODAMA to multiple spatial transcriptomics datasets generated using diverse technologies, including MERFISH, 10x Visium, and Visium HD. KODAMA effectively recovered expected anatomical structures, such as hypothalamic subregions and cortical layers (**Fig. 2** and **3**), and outperformed non-spatial methods in preserving spatial continuity. In addition to known features, KODAMA identified biologically relevant patterns missed by other algorithms, including frostbite artifacts and sublayer stratification in the DLPFC dataset (**Fig. 3**). Quantitative evaluation consistently demonstrated KODAMA’s superior performance compared to other dimensionality reduction and clustering tools. In addition, we applied the KODAMA framework to identify spatially resolved genetic alterations within neoplastic regions of prostate tissue (**Fig. 4**) and colorectal carcinoma (**Fig. 5**), revealing gene expression gradients and tumor-specific molecular patterns."

**Fig. 2:**
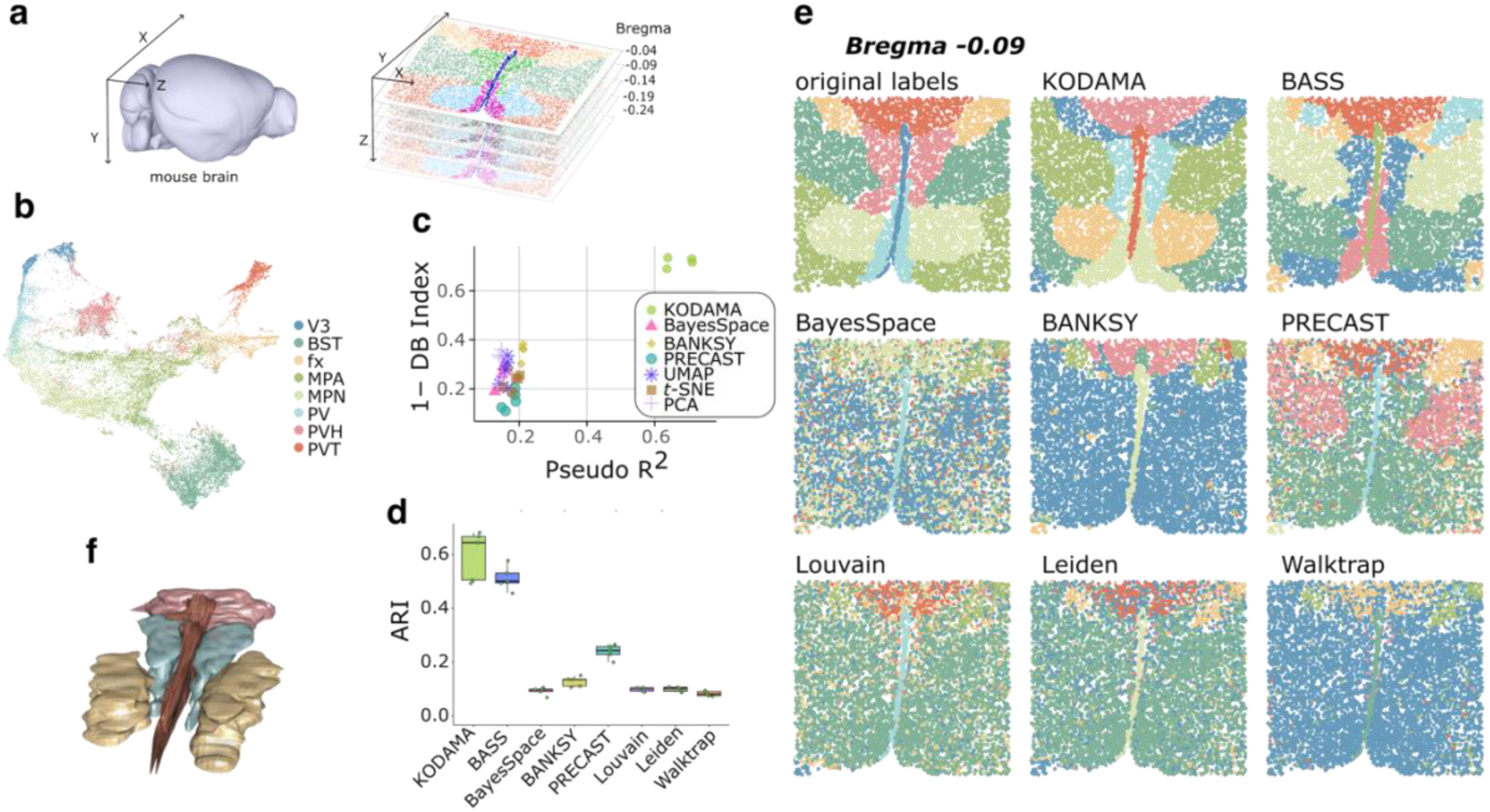
MERFISH dataset. **a,** A 3D model of the mouse brain, adapted from the Allen Mouse Brain Common Coordinate Framework (CCFv3) [43], with main anatomical axes (X, Y, Z). On the right, is a view of spatial slices showing the distribution of transcriptomic points at different Bregma levels. **b)** KODAMA spatial dimension colored based on the different tissue microarchitectures, including the third ventricle (V3), bed nuclei of the stria terminalis (BST), columns of the fornix (fx), medial preoptic area (MPA), medial preoptic nucleus (MPN), periventricular hypothalamic nucleus (PV), paraventricular hypothalamic nucleus (PVH), and paraventricular nucleus of the thalamus (PVT). **c)** Comparison of tested feature extraction methods using 1-DB-Index and Pseudo-R². **d)** Boxplot of the ARI comparing the accuracy of different clustering algorithms (KODAMA, BayesSpace, BANKSY, PRECAST, Louvain, Leiden, and Walktrap) against the ground truth annotations. **e)** Clustering results for a slice at Bregma −0.09 mm. The compared methods include KODAMA, BASS, BayesSpace, BANKSY, PRECAST, and three classical clustering approaches (Louvain, Leiden, Walktrap). Ground truth annotations (Real Labels) are also included as a reference. **f)** 3D reconstruction of spatial region domains identified by KODAMA within an anatomical structure of the mouse brain.

**Fig. 3:**
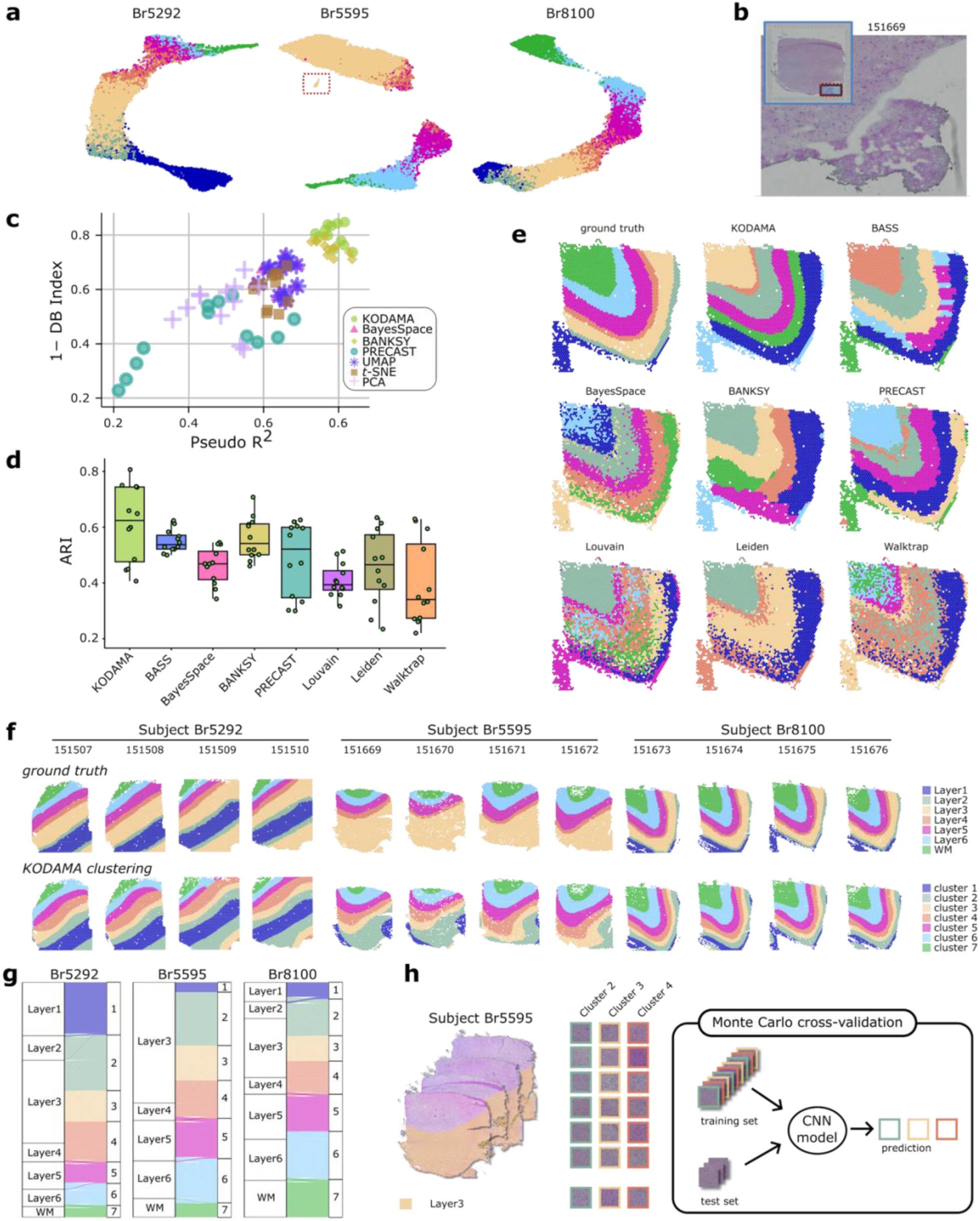
DLPFC dataset. **a,** KODAMA output for each subject analyzed colored by ground truth assignment. **b,** H&E histological image identifying the frozen bite region of the sample 151669. **c,** Comparison of tested feature extraction methods using 1-DB-Index and Pseudo-R². **d,** Boxplot of the ARI comparing the accuracy of different clustering algorithms. **e,** Clustering results for slide 151675. **f,** KODAMA clustering on the integrated 12 DLPFC samples reveals new sublayers for the subject Br5595. **g,** Sankey diagram of the comparison between the ground truth layers and the clustering results of the subject Br5595 obtained on KODAMA performed on all 12 slides. **h,** Workflow of the deep learning analysis performed to evaluate the predictivity of sublayer classification using the H&E image tiles.

**Fig. 4:**
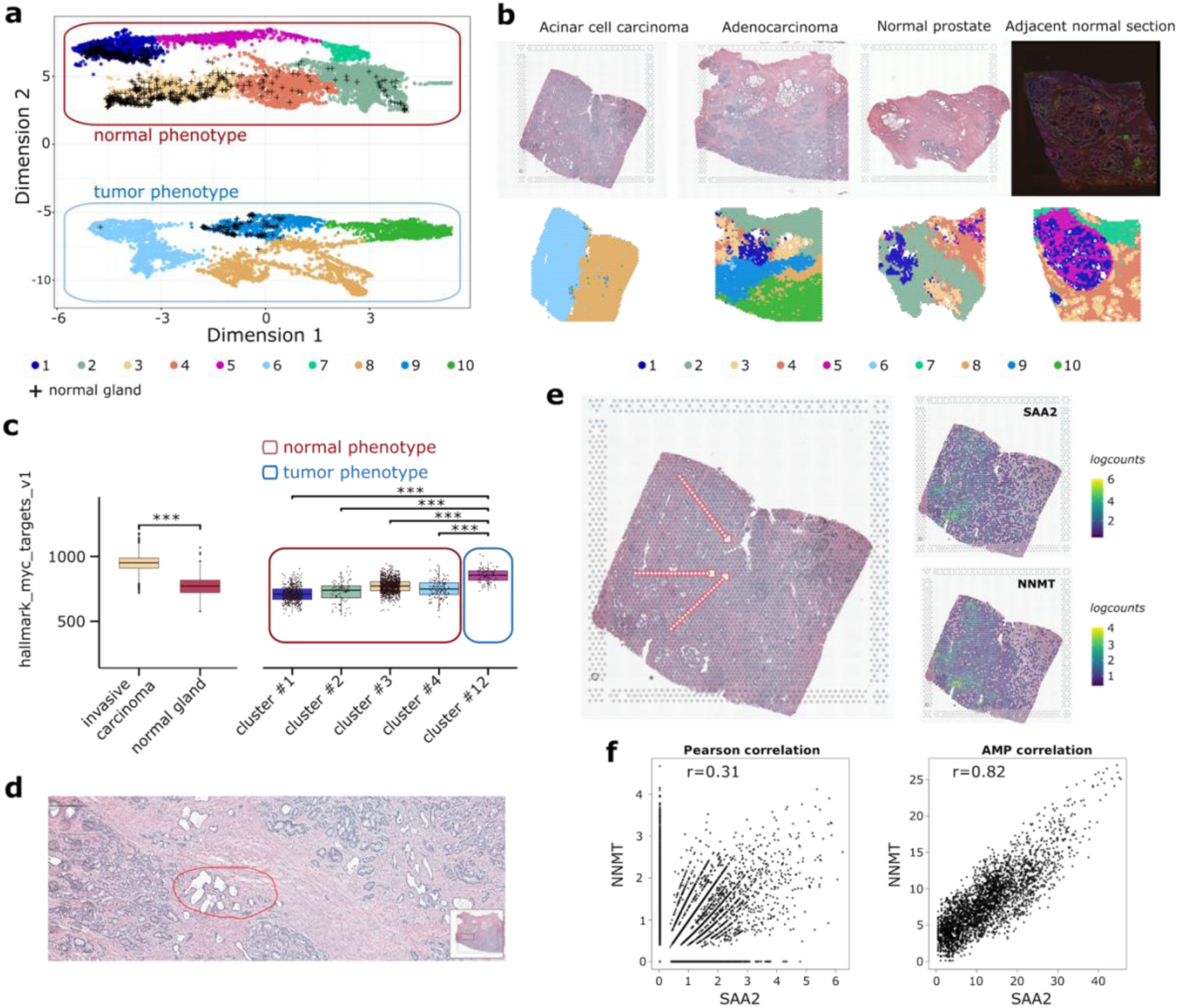
Prostate tissue dataset. **a,** KODAMA visualization of the four prostatic tissues color-labeled by clustering. **b,** Histological H&E staining of the four 10x Visium prostate tissue samples and clustering from the KODAMA output. **c,** MYC gene set enrichment value across the pathological annotations and the KODAMA clustering. **d,** Identification of a tissue region labeled as normal on QuPath. **e,** Trajectory analysis showing the variations in gene expression levels. **f,** Comparison of Pearson and AMP correlation test showing the spatial gene-gene correlation.

**Figure 5:**
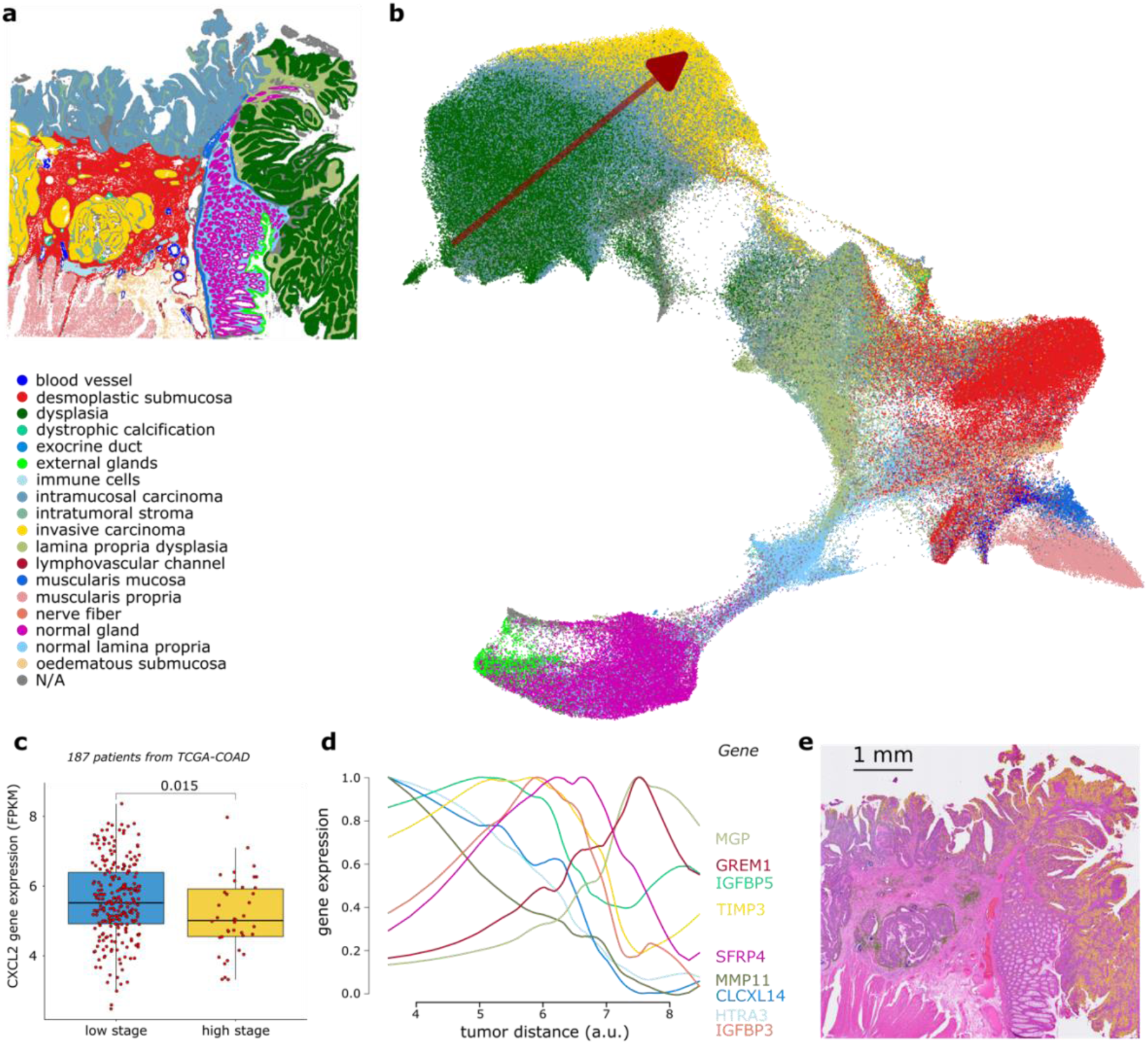
Spatial analysis of a VisiumHD colon cancer dataset using KODAMA. **a,** Pathological annotation of the tissue. **b,** KODAMA output showing the change in gene expression profile from dysplasia to intramucosal carcinoma and invasive cancer regions. **c,** Comparison of CXCL2 gene expression in bulk RNA profile between low and high stage colon carcinoma in TCGA-COAD cohort. **d,** Evolution of the expression of multiple genes as a function of distance from the cancer. **e,** Histological image illustrating the spatial distribution of specific genes (*i.e.*, MMP11 in green and CXCL2 in yellow) in different tissue regions, indicating potential roles in tumor transition.

### Capturing Anatomical Structures in 3D Spatial Single-Cell Data

The structure-function relationship in tissue microanatomical structures is particularly evident in the laminar organization of the human cerebral cortex, where cells in different cortical layers exhibit distinct gene expression profiles, as well as varied patterns of morphology, physiology, and connectivity. Here, we used a dataset of 155 genes generated from the hypothalamic preoptic region of a healthy mouse using MERFISH technology [40]. The data were filtered to include samples from five consecutive brain sections from the hypothalamic regions: Bregma −0.04 mm, −0.09 mm, Bregma −0.14 mm, Bregma −0.19 mm, and Bregma −0.24 mm (**Fig. 2a**). The Bregma is a landmark point where the coronal and sagittal sutures of the skull intersect, and brain coordinates are measured relative to this point in millimeters. On average, each image contains 5,663 cells distributed across eight microanatomical regions [40].

We applied KODAMA to the full set of 28,317 cells, incorporating their three-dimensional spatial coordinates. By integrating spatial information directly into the KODAMA algorithm, we achieved accurate delineation of microanatomical tissue structures within a two-dimensional projection space (**Fig. 2b**). To quantitatively assess the ability of each dimensionality reduction method to resolve true microanatomical compartments, we employed the McFadden-adjusted pseudo-R² [41] as a measure of fidelity to represent the ground truth annotations, and the inverse Davis–Bouldin Index (1 – DB Index) [42] as a proxy for cluster compactness and separation. We benchmarked KODAMA against established spatial and non-spatial feature extraction approaches, including BayesSpace, BANKSY, PRECAST, UMAP, *t*-SNE, and principal component analysis (PCA). Among all the evaluated methods, KODAMA consistently yielded the highest pseudo-R² and 1 – DB Index values, indicating superior performance in preserving both anatomical fidelity and cluster coherence (**Fig. 2c**). The outputs of all compared methods are visualized in **Supplementary Fig. 1**.

Conventional PCA, used for dimensionality reduction of this dataset, failed to adequately resolve distinct tissue compartments, leading to ambiguous and difficult-to-interpret projections. We demonstrate that augmenting PCA with the message-passing preprocessing markedly enhances its discriminatory power, particularly within the first two principal components (**Supplementary Fig. 2**). Message-passing preprocessing similarly improved the performance of other non-spatial techniques, such as *t*-SNE and UMAP, as evidenced by both visual separation (**Supplementary Fig. 2**) and quantitative gains in pseudo-R² and 1 – DB Index (**Supplementary Fig. 3**).

To cluster the different spatial region domains, we applied the Louvain algorithm to the KODAMA’s two-dimensional projection space and subsequently refined the clustering result using the SVM model. Clustering accuracy was assessed using the Adjusted Rand Index (ARI) across multiple algorithms, including BayesSpace, BANKSY, PRECAST, Louvain, Leiden, and Walktrap (**Fig. 2d)**. KODAMA consistently outperformed other methods, achieving higher alignment with ground truth annotations. A focused comparison of clustering results was performed on a spatial slice at Bregma −0.09 mm, involving advanced methods (KODAMA, BASS, BayesSpace, BANKSY, PRECAST) and non-spatial clustering techniques (Louvain, Leiden, Walktrap) (**Fig. 2e)**. The spatial mappings of the clustering results are visualized in **Supplementary Fig. 4**. KODAMA exhibited superior spatial continuity and biological relevance, closely matching known anatomical annotations, highlighting its robustness in handling complex spatial data.

To evaluate independently the tools integrated into the KODAMA framework, we tested message-passing preprocessing and SVM refinement of the non-spatial clustering techniques (*i.e.*, Louvain, Leiden, and Walktrap). These tools enhance the identification of tissue segments in non-spatial clustering techniques in terms of ARI (**Supplementary Fig. 5**). The spatial mappings of the clustering results is visualized in **Supplementary Fig. 6**.

We extended our framework to include a function for interactive 3D visualization of the identified clusters within their anatomical context. (**Fig. 2f**). This interactive visualization provides an intuitive way to explore spatial relationships and may help to reveal biologically meaningful insights into spatial interactions between neighboring regions.

### Identification of novel tissue segments in multi-slide DLPFC datasets

Understanding the spatial organization of gene expression within the brain is essential for uncovering region-specific mechanisms underlying neurological disorders. ST provides a powerful approach to map transcriptional activity directly onto tissue architecture, enabling the identification of molecular signatures associated with defined anatomical regions. We applied KODAMA to examine data from the human dorsolateral prefrontal cortex (DLPFC) obtained using the 10x Visium platform [44]. The dataset contains expression values of 33,538 genes measured on two pairs of tissue sections from three independent neurotypical adult donors. Each pair consists of two directly adjacent, 10 μm serial tissue sections, with the second pair located 300 μm posterior to the first, resulting in a total of 12 tissue sections. Each subject contributed four slices, identified as 151507, 151508, 151509, and 151510 for first subject Br5292; 151669, 151670, 151671, and 151672 for second subject Br5595; and 151673, 151674, 151675, and 151676 for third subject Br8100. Each sample was manually annotated [44] to define its cortical layers, resulting in either five or seven layers per sample, including the white matter (WM) as one of the annotated regions. Processed subject data comprises 17,734, 14,646, and 13,938 spot counts for subjects Br5292, Br5595, and Br8100, respectively. We did not report a relevant batch effect in the gene expression between sections from the same donor. To compare different methods independently by the batch effect correction algorithms, such as Harmony [45], we performed subsequent analyses separately on the four tissue slides of each subject.

KODAMA analysis highlighted underlying relationships between datapoints that corresponded with different functional regions (**Fig. 3a**). The results indicated a continuity between each layer rather than a marked separation. Interestingly, KODAMA also detected a small subcluster of datapoints in the subject Br5595, which corresponds to a section of tissue damaged during freezing (**Fig. 3b**) as observed by histological analysis. Overall, KODAMA emerged as the best feature extraction method in terms of pseudo-R² metric and DB-Index, followed by BANKSY (**Fig. 3c**). Non-spatial informed approaches like PCA, UMAP, and *t*-SNE exhibited inferior performance due to their inability to exploit the spatial structure of the data. The results of each feature extraction method are shown in **Supplementary Fig. 7**. We next clustered the different spatial regions on the KODAMA dimensional space using the Walktrap method and we refined the clustering results using SVM. Our results showed a higher ARI index compared to other methods (**Fig. 3d**. A visual investigation of the results of each method reveals that despite the high value of ARI, some methods do not show an optimal stratification of the layers (**Supplementary Fig. 8**). In **Fig. 3e**, we showed the clustering results for the data sample 151675.

The simultaneous analyses of all twelve patients slides provided additional insight into the biology of these tissues (**Supplementary Fig. 9**). For example, an outlier cluster in the KODAMA output was spotted and associated with a higher mitochondrial percentage in the slide belonging to the subject Br5595 (**Supplementary Fig. 10**). As suggested in [46], the higher mitochondrial gene ratio could mark the high focal stress point of excitatory neurons. After removing these datapoints, we clustered the KODAMA output using k-means clustering, identifying a sub-segmentation of layer 3 (**Fig. 3f**). This stratification also appears using a novel rainbow (a.k.a. RGB) visualization approach that improves the colors’ separation. **Supplementary Fig. 11** shows the color association of each spot in the KODAMA dimensionality space and slides. The Sankey diagram **(Fig. 3g)** illustrates correspondences between layers, offering an intuitive representation of relationships between spatial layers and the additional identified groups in each subject.

Next, we analyzed the differentially expressed genes in the identified sublayers (*i.e.*, clusters 2, 3, and 4) of the Layer 3 (**Supplementary Table 1**), revealing distinct physiological roles. *MALAT1*, *PPP3CA*, and *CAMK2N1*, enriched in cluster 2, are involved in brain function, particularly related to calcium signaling and synaptic plasticity. *PPP3CA* encodes calcineurin, a protein phosphatase crucial for calcium signaling [47], while *CAMK2N1* encodes a protein kinase inhibitor that regulates calcium levels [48]. Although it has been shown that *MALAT1* level modulates synapse density in cultured neurons [49], its role in calcium-related mechanisms has been explored only in unrelated fields [50]. Cluster 3, marked by *CCK*, *RGS4*, and *CALM2*, reflects neurotransmitter signaling and interneuron modulation [51]. Their downregulation in patients with Alzheimer’s disease [46] could suggest the importance of this sublayer classification in the understanding of this disease. Cluster 4, dominated by *MBP*, *PLP1*, and *CNP*, is specialized in myelination and axonal integrity [52]. These profiles highlight a transition from synaptic processing to structural and conductive support. This level of resolution provided a more detailed perspective on the molecular architecture of the cortex, uncovering subtle transcriptional differences that contribute to cortical organization.

To further validate the identified sublayers, we investigated the possibility of distinguishing the sublayers by digital analysis of the histological H&E images (**Fig. 3h**). A set of histological image tiles (∼ 270μm×270μm size) centered in the Visium spots from layer 3 of the subject “Br5595” was extracted and associated with previously identified sublayers. The sublayers 1, 3, and 5 contained, respectively, 3610, 1022, and 2210 Visium spots. To have a relevant number of images layers 3 and 5 were combined. Then, we built a Monte Carlo (MC) cross-validation procedure to estimate the prediction accuracy of the convolutional neural network (CNN) model built on a balanced training set comprising 75% of all tiles. The sublayers identified by KODAMA were predicted with an average accuracy of 66.9±5.7%. This clustering provides novel insights into the spatial organization of the DLPFC and highlights KODAMA’s ability to detect subtle variations in tissue architecture.

### Deciphering the molecular landscape of prostate tissue

Tumor heterogeneity results in the high complexity and diversity of malignant tumors during their evolution, while drug resistance caused by tumor heterogeneity is a significant challenge in current cancer research and antitumor drug discovery [53, 54]. ST offers a significant advantage by facilitating comprehension of the spatial arrangement of diverse cell types within the tumor microenvironment and furnishing crucial spatial details concerning the transcriptomic landscape of tumor cells. We analyzed a prostate tissue dataset, including normal prostate, two adenocarcinomas, and adjacent normal sections retrieved from the 10X Visium database. The ground truth for an adenocarcinoma tissue was obtained from the 10X website, while the pathological annotation for the other tissues was provided by two pathologists (B.P. and F.S.) and then imported using the SpatialData framework [55]. The four tissues were integrated using Harmony, and PCA was employed to visualize the variability both before and after integration (**Supplementary Fig. 13**). KODAMA effectively distinguished different tissue types, revealing notable differences between healthy and cancerous regions (**Supplementary Fig. 14**). The integration of data across multiple samples allowed for a more robust identification of shared features and unique patterns. KODAMA’s clustering identified key regions of benign prostatic hyperplasia (BPH), including BPH glands (cluster 1) and BPH stroma (cluster 5), present in both malignant and adjacent normal tissues, highlighting the spatial organization of these structures (**Fig. 4a-b**) as evident in both normal tissue sections.

In the adenocarcinoma slide, KODAMA analysis uncovered the subtle pathological changes in normal glands. Some glands adjacent to tumor tissues indicated as normal showed to have a phenotype close to the tumor. Gene set enrichment analysis illustrated elevated MYC expression in these normal glands, suggesting that even seemingly healthy regions may harbor pre-cancerous characteristics (**Fig. 4c** and **Supplementary Table 2**). With the KODAMA framework, we included a function to export selected tissue regions in a GeoJSON file format for microscopy software. To a deeper histological evaluation, these glands could be referred as prostatic intraepithelial neoplasia (PIN), as shown in **Fig. 4d**. These regions within glandular lining cells, indeed, exhibit cytological abnormalities, including nuclear overlapping, nuclear hyperchromasia, irregular nuclear borders, and prominent nucleoli, while remaining within the existing duct structure. These features resemble cancer cells without fibromuscular stromal invasion.

In the second slide of adenocarcinoma (named by 10X as acinar cell carcinoma), KODAMA clustering delineated the gene expression differences across three prostatic lobes (**Fig. 4b**). We built a user-guided trajectory that allows us to trace the progression of cancer from the peripheral regions toward the central core of the tissue (**Fig. 4e**), offering insights into the tumor’s developmental pathway. The correlation between gene expression and trajectory is shown in **Supplementary Table 3**. We identified Serum Amyloid A2 (*SSA2*) and Nicotinamide N-methyltransferase (*NNMT*) overexpressed mainly in the peripheral regions of the prostate (**Fig. 4e**). *NNMT* is highly expressed in various human cancers and is associated with proliferation, invasion, and metastasis [56–58]. In bladder cancer, the elevated *NNMT* expression within cancer-associated fibroblasts drives the overexpression of serum amyloid A family proteins and fosters an immunosuppressive tumor microenvironment by recruiting and modulating macrophages, ultimately facilitating tumor progression and resistance to immunotherapy [59].

To investigate the potential interrelationship between *SSA2* and *NNMT* expression in prostate cancer, we applied Pearson’s correlation to the data. To address the sparse and discontinuous ST data, where missing values disrupt the detection of biologically relevant relationships, we applied after message-passing procedure leveraging spatial locality (**Fig. 4f**). This substantially improved the resulting correlation coefficient, indicating a strong interaction between these genes, addressing key limitations in conventional correlation analysis. Additional studies in ovarian cancer indicate that *NNMT*-expressing fibroblasts secrete 1-methylnicotinamide (MNA) – an immunoregulatory metabolite capable of shaping T cell function – which further highlights *NNMT*’s capacity to modulate anti-tumor immunity [60]. Although direct evidence in prostate cancer remains limited, the parallels in fibroblast-driven immunomodulation point toward a potential role for *NNMT* in prostate tumorigenesis.

### Assessment of handling large datasets (Visium HD)

The VisiumHD platform, recently introduced by 10X Genomics, has reduced the diameter of spots from 55 μm to an edge length of 2 μm, eliminating gaps between spots. The significant enhancement enables gap-free and bias-free single-cell resolution. VisiumHD slides contain two 6.5×6.5 mm capture areas, each consisting of ∼11 million 2×2 µm squares arranged in a continuous array of uniquely barcoded oligonucleotides. To evaluate its capability in handling large datasets, we applied KODAMA to VisiumHD data acquired from human colon cancer slide at a resolution of 8×8 µm squares. The histological H&E image of colorectal tumor tissue was annotated by two pathologists (B.P. and F.S.) using the QuPath software [39] and subsequently imported into our framework using SpatialData (**Fig. 5a**). KODAMA clearly identified distinct clusters representing different tumor microenvironments (**Fig. 5b**). These clusters represent histologically relevant regions, including invasive carcinoma, dysplastic epithelium, and desmoplastic stroma, which are critical for understanding tumor heterogeneity.

KODAMA results highlighted the gradient changes between dysplasia and invasive carcinoma. To retrieve the KODAMA gradient associated with each spot, we developed a trajectory analysis to focus specifically on a section of KODAMA results (**Fig. 5b**). We ranked a list of genes associated with the KODAMA gradient (**Supplementary Table 4**). We identified CXCL2, a pro-inflammatory chemokine involved in regulating the tumor microenvironment, as strongly associated with cancer progression. CXCL2 is less expressed in invasive cancer.

The decrease in CXCL2 expression in the invasive zone (**Supplementary Fig. 15**) may suggest a loss of inflammatory activity at this advanced stage, indicating an adaptation of the tumor microenvironment toward a more permissive state for invasion [61]. This gene plays a crucial role in recruiting neutrophils and macrophages to the tumor microenvironment, suggesting an immune escape of the invasive carcinoma [62]. To validate CXCL2 as a marker of tumor progression, we examined its expression value in the TCGA-COAD cohort (280 patients). Our analysis revealed a downregulation of CXCL2 in patients with high (*i.e.*, T4 stage) tumor stage (**Fig. 5c**). These findings provide interesting insights into tumor microenvironment remodeling and suggest that certain chemokines could represent potential therapeutic targets to limit colorectal cancer progression [63].

Additionally, we investigated gene gradients in proximity to invasive carcinoma. We identified a list of genes whose gene expression correlates with the distance from the invasive carcinoma using the Maximal Information Coefficient (MIC) as a score (**Supplementary Table 5**). **Fig. 5d** shows a panel of genes (with MIC>0.8), including IGFBP5, Matrix Gla Protein (MGP), Tissue Inhibitor of Metalloproteinases 3 (TIMP3), HtrA serine peptidase 3 (HTRA3), and matrix metalloproteinases 11 (MMP11), with a different gradient across the desmoplastic submucosa (**Supplementary Fig. 16-20**), suggesting a deregulation of the proteases’ activities. MMP11 is highly expressed around the invasion zone, suggesting its potential role in tumor transition, while CXCL2 is activated in dysplastic regions, indicating early regulation in tumor progression. MMP11 and CXCL2 are visualized in **Fig. 5e**. These results suggest that the tumor microenvironment actively remodels the extracellular matrix and alters the balance of pro- and anti-invasive signals [63].

## Discussion

Spatially informed KODAMA is a powerful and innovative solution for ST analysis. By integrating spatial constraints directly into its algorithm for feature extraction, crucial spatial relationships between cells are preserved, which is essential for considering cellular interactions and microenvironmental dynamics. In our previous implementation of KODAMA, we introduced the concept that the separation between samples can emerge in a vector of labels where these are swapped to achieve a high cross-validation accuracy. This implementation has already been successfully used to profile single-cell data sets [64]. ST data analysis is often challenging due to the inherent sparsity of sequencing coverage, where many genes exhibit low or undetected expression across spatial locations due to technical limitations, sequencing depth, or biological variability. Additionally, noise arising from cell-to-cell variation further complicates the analysis, making it crucial to employ robust computational methods for accurate interpretation. Swapping the labels of samples that are spatially close provides KODAMA with guidance toward maximizing cross-validation accuracy based on spatial relationships rather than relying solely on feature similarity. This spatially informed adaptation of KODAMA ensures that local spatial structures are effectively leveraged, significantly enhancing the interpretation of ST data.

While we primarily focused on KODAMA’s dimensionality reduction capabilities, KODAMA dimensions can be used as input for clustering methods to identify tissue domains. KODAMA was applied to a MERFISH data set highlighting its capacity to handle 3D spatial data and to be used for the analysis of large collections *i*ST datasets. The introduction of several new tools extends KODAMA into a comprehensive framework for ST analysis. This includes the message-passing pre-processing procedure that combines weighted phenotypic information across spatially close cells, emphasizing a distinct fingerprint in each tissue section. The message-passing approach has also demonstrated benefits for other methods that do not explicitly incorporate spatial information. After the clustering step, the KODAMA framework includes a tool that sharpens tissue segment boundaries, potentially increasing spatial resolution. We demonstrate that applying SVM-based refinement effectively improves tissue segmentation quality across datasets with varying proportions of cell populations, such as MERFISH data, and datasets featuring laminar tissue architectures, exemplified by the DLPFC.

Clustering methods often rely on assumptions about data distribution and cluster shapes [65], which may not hold for all tissue types, leading to less accurate results. Traditional clustering approaches also fail to capture continuous spatial trends, as seen in the DLPFC dataset. Instead of distinct clusters, KODAMA permits the observation of a transcription gradient across different regions, highlighting the limitations of standard clustering techniques in preserving spatial continuity. We suggest investigating the results of KODAMA in detail before deploying any clustering algorithm. Further, KODAMA was also able to detect the presence of regions with altered gene expression due to sample damage through freezing.

KODAMA led to the identification of a sub-clustering tissue segmentation that emerged when all 12 tissues were integrated using leveraging. Differential gene expression analysis points out putative markers and enriched pathways in these sublayers, and, for the first time, the clustering obtained from the KODAMA analysis has been validated using a CNN model built on the histological H&E image of the tissue.

In prostate tissue, KODAMA identified spatially heterogeneous gene expression across normal glands, unveiling tissue regions with cytological abnormalities. The KODAMA framework gives the possibility to identify gene changes across a trajectory drawn on the histological image. This allows us to identify genes overexpressed in the exterior area of the prostate. Standard methods like Pearson’s correlation often struggle with sparse and discontinuous ST data, where missing values disrupt the detection of biologically relevant relationships. Here, we introduced a new powerful approach to deciphering gene-gene interactions by leveraging spatial locality, addressing key limitations in conventional correlation analyses.

With the design of the landmark approach and the possibility to run in parallel the iterative procedures of cross-validated accuracy maximization implemented in this new version of KODAMA, we pave the way to the analysis of data sets composed of hundreds of thousands (possibly millions) observations.

KODAMA has been successfully integrated with established ST frameworks such as SpatialExperiment, Seurat, and Giotto. The findings underscore KODAMA’s broader applicability in ST, where maintaining spatial continuity is crucial for biological interpretation. The method’s ability to integrate complex spatial data without imposing arbitrary boundaries enhances its utility in studying tissue heterogeneity, microenvironmental interactions, and potential biomarkers. By leveraging this approach, previously overlooked molecular patterns can be revealed, contributing to a deeper understanding of cancer progression and potential therapeutic interventions.

## Conclusion

KODAMA’s computational efficiency allowed for the analysis of benchmarking datasets that presented multiple challenges, including diverse data types (array-based and image-based platforms), multi-slide integration, complex clustering patterns, and high-dimensional spatial trends. By addressing these challenges, KODAMA preserved the biological context of spatial data while enabling the identification of tissue-specific cellular clusters and gene expression gradients. This scalability made it possible to process large datasets efficiently while maintaining the spatial structure necessary for meaningful biological interpretation.

## Methods

### KODAMA framework

#### Overview of the spatially informed KODAMA

The KODAMA algorithm comprises five key steps: initialization, iterative refinement, prediction of projection observations, dissimilarity matrix calculation, and dimensionality reduction. The iterative refinement procedure, encompassing the initial three steps, inherently produces suboptimal solutions due to its stochastic nature; therefore, multiple iterations (100 by default) are performed to average these random effects.

In the initialization step, the algorithm selects a representative subset of samples, known as landmark observations, on which the next step will be performed. These landmarks are chosen to capture the full diversity within the dataset. If the number of landmarks is not explicitly specified, the default is set at 75% of the total observations. Consequently, the dataset is partitioned into two subsets: the landmark set and the projection set, consisting of remaining observations.

To ensure landmark observations accurately reflect the dataset’s heterogeneity, K-means clustering is employed across the feature space. Initially, the data is segmented into clusters matching the predefined number of landmarks, and a single random observation from each cluster is selected as a landmark. Following landmark selection, an initial vector of classification labels is assigned by grouping observations based on feature similarity into clusters defined by a parameter called *splitting*.

When spatial information is incorporated, an additional clustering procedure is performed based on spatial dimensions. The initial number of spatial clusters is determined by a *resolution* parameter (default value: 0.3), calculated as approximately the product of the total number of observations and the *resolution* value. Subsequently, the labels assigned to the landmark observations inside the same spatial cluster are forced to be assigned to the same class.

The second step involves iteratively refining the classification labels assigned to landmark observations. This step aims to maximize cross-validated accuracy using a supervised learning approach previously described [31]. Specifically, Partial Least Squares (PLS) discriminant analysis serves as the supervised model for calculating cross-validated accuracy, facilitated by our new implementation of Partial Least Squares (PLS) regression. Throughout each iteration, randomly selected misclassified observations have their class labels swapped to match the predictions, and a new calculation of cross-validated accuracy is performed. If spatial information is provided, observations within the same spatial cluster are forced to share the same class label during the iterative process. This strategy preserves local spatial relationships during classification refinement. The procedure is guided by choosing the classification vector with the best cross-validated accuracy during each iteration.

To improve computational efficiency, the algorithm has been adapted for parallel processing, distributing tasks across multiple computational cores. This parallel implementation significantly decreases computation time and optimizes resource utilization. By combining landmark selection and parallel computing, the KODAMA algorithm demonstrates scalability suitable for handling complex, large-scale transcriptomics datasets without compromising accuracy or responsiveness.

This iterative refinement is exclusively performed on landmark samples. In the third step, the optimized classification labels obtained from this procedure are used to train a PLS-DA model on the landmark observations, allowing the classification of projection samples. When spatial information is included, all observations—both landmarks and projections—within the same spatial cluster are constrained to share identical class labels. This ensures spatial coherence in the classification results. In the fourth step, a partial dissimilarity analysis is performed by determining the indices and calculating the KODAMA distances of the closest observations. Specifically, the number of nearest observations considered is calculated as one plus the minimum among the following three values: the number of landmarks, one-third of the total number of observations, and 500. Constructing this partial dissimilarity matrix helps to mitigate memory complexity issues, as memory requirements for a full dissimilarity matrix scale quadratically with the number of observations. However, calculating a complete square dissimilarity matrix remains feasible for smaller datasets.

In the fifth and final step, the dissimilarity matrix is converted into a reduced number of KODAMA dimensions (two by default). This dimensionality reduction is typically performed using algorithms such as UMAP (default) or *t*-SNE. Alternatively, multidimensional scaling may be applied if a complete square dissimilarity matrix is available.

#### Message-passing

This method aggregates phenotypic information from neighboring cells or spots, weighted inversely by their spatial distance.

In mathematical terms, the message-passing approach transforms feature values according to the following formula:

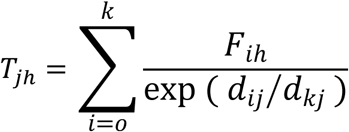

where *T_jh_* is the transformed value of the feature *h* of the focal cell *j*; *k* (default: 15) is the chosen number of nearest neighbors (NN) to the cell *j*; *d_ij_* denotes the distance between the focal cell *j* and neighbor cell *i*, with *i=0* corresponding to the focal cell itself (thus, *d_ij_=0*); *d_kj_* indicates the distance from the focal cell to its furthest neighbor among the *k* nearest neighbors; and *F_ih_* is the original feature value *h* in cell *i*.

The rationale behind this approach is to ensure that each cell incorporates spatially weighted features from its local neighborhood. This allows for capturing both phenotypic similarity and spatial proximity, effectively preserving spatial coherence within the dataset. This method particularly benefits spatial transcriptomics studies by enhancing the identification of biologically relevant tissue microstructures and gradients, which might otherwise remain undetected due to local noise or spatial discontinuities.

#### After Message-Passing (AMP) correlation

This preprocessing step leads to a novel statistical method designed specifically to compute correlations in spatially resolved transcriptomics data. ST data typically exhibit sparse expression profiles, characterized by low detection rates and considerable numbers of missing gene expression values due to limitations in the sensitivity of current technologies. Traditional correlation coefficients, such as Pearson or Spearman, often fail to adequately capture meaningful relationships between genes under these sparse conditions, since the low detection rate can obscure underlying biological associations.

The AMP preprocessing mitigates this challenge by leveraging spatial information to reconstruct gene expression signals within local neighborhoods. By aggregating gene expression data from nearby cells or spots, the method effectively compensates for missing or low-intensity signals, thus enhancing the robustness and interpretability of subsequent correlation analyses. Specifically, AMP exploits spatial continuity, using weighted averaging based on proximity, thereby inferring missing information from spatially correlated observations. This allows for more accurate and biologically relevant correlation estimates, uncovering subtle yet significant patterns of gene co-expression that would otherwise remain undetected. Consequently, the AMP correlation method represents an advancement over traditional correlation measures by integrating spatial locality directly into statistical inference, enabling a deeper understanding of the spatial dynamics underlying gene interactions within tissue microenvironments.

#### Support Vector Machine (SVM) refining

To accurately identify tissue microarchitectures based on spatial transcriptomics data, a refinement procedure leveraging SVM, a robust supervised learning algorithm, is employed. Following a clustering procedure, cluster boundaries are further refined using SVM classification. Specifically, the initial clustering results, combined with their corresponding spatial coordinates, serve as the training dataset for the SVM classifier.

The SVM model, utilizing a radial kernel, determines optimal decision boundaries by maximizing margins between predefined clusters based on spatial coordinates. By computing planes (in three-dimensional space) or lines (in two-dimensional space) within the spatial domain, SVM effectively delineates cluster boundaries with enhanced precision. These SVM-derived decision boundaries are then applied to reclassify spatial observations, improving the clarity and consistency of tissue cluster assignments.

The refinement process involves partitioning the spatial data into smaller subsets (tiles), within which local decision boundaries are independently optimized through separate SVM models. When additional spatial observations at higher resolution are provided, the trained SVM models predict their cluster memberships based on the previously established spatial decision boundaries.

The refined cluster boundaries are subsequently projected back onto the original spatial domain, significantly enhancing interpretability. This tile-based, SVM-driven refinement approach is specifically designed for spatially informed datasets, effectively improving spatial coherence and delivering biologically meaningful delineations of microarchitectural features within the analyzed tissues.

#### Exporting spatial geometry object

Selected observation coordinates (*e.g.*, Visium spots belonging to a cluster) can be converted into GeoJSON polygons to enable pathological analysis using software such as QuPath. Spatial clusters derived from these selected coordinates are converted into concave polygon boundaries using a concave hull algorithm [66] implemented in the concaveman R package. Each polygon boundary is then smoothed using Gaussian kernel smoothing (smoothing factor = 3) implemented in the R package ‘smoothr’, refining polygon boundaries and ensuring biological relevance and visual clarity. Finally, these smoothed polygons were exported as GeoJSON files using the sf R package, facilitating their subsequent visualization and integrated pathological analysis in QuPath.

#### Integration with different pipelines

KODAMA is implemented as a flexible and interoperable algorithm that is fully compatible with standard R data structures, specifically leveraging the S3 method system for generic function compatibility. Input datasets for KODAMA are standard R matrices, ensuring broad usability across diverse analytical workflows, not limited to spatial transcriptomics. The structured implementation of S3 methods within KODAMA ensures seamless integration with widely used bioinformatics frameworks such as Giotto, Seurat, and SpatialExperiment, as well as other R-based analytical ecosystems. Additionally, the KODAMA algorithm efficiently handles both sparse and dense matrix formats, directly accepting these formats without requiring specialized preprocessing. Consequently, this robust compatibility simplifies incorporation into existing analytical pipelines, enabling accessible and efficient deployment for high-dimensional biological data analysis across multiple applications.

#### User-guided trajectory

This R function constructs a smoothed trajectory from a set of two-dimensional coordinates and optionally computes mean values along that trajectory. It begins by checking whether the user has already supplied a trace of points, or else uses the identify function to interactively select ordered points on a plotted plane. After sorting these control points, the function applies xspline to generate a smooth curve through them, then refines the output by interpolating at regular intervals. This yields a denser sequence of (x, y) coordinates, from which cumulative distances are calculated to resample the curve at a user-specified resolution. If a separate data matrix is provided, each trajectory coordinate is matched to its k nearest neighbors in the original input space, and the mean values of those neighbors are computed. The function plots the resulting spline curve on the existing figure, marking the first point of the trajectory distinctly from the others, and finally returns a list containing the smoothed curve’s coordinates, the mean values of the data across nearest neighbors (when applicable), the indices of those nearest neighbors, and relevant settings used in the process.

#### Trajectory RGB visualization

A trajectory is inferred using the slingshot function from the slingshot R package [67] applied to the dimensionally reduced data (*i.e.*, KODAMA dimensions). To assign trajectory-based colors to each observation, a rainbow color palette was generated corresponding to points evenly distributed along the inferred trajectory. Each observation was mapped to the nearest trajectory point using NN search using the R package Rnanoflann, inheriting its respective color assignment. Observations were visualized as colored points representing their relative position along the trajectory. The trajectory itself was overlaid, highlighted by distinct markers to enhance the interpretability and clarity of the inferred progression patterns.

#### Spatially variable genes

Spatially variable genes were identified using the SPARK-X algorithm [68]. SPARK-X employs efficient statistical modeling and hypothesis testing to rapidly detect these spatially variable genes, even in large-scale datasets, helping researchers uncover genes linked to spatially organized biological processes and tissue architecture. The genes are ranked based on the p-values. In the case of multiple samples analysis, SPARK-X is performed independently for each sample and the genes are ranked based on the minimum value across all samples

### Performance evaluation

Different metrics were used to evaluate the performance of the methods tested. ARI [69] was used to measure the goodness of the different clustering results. ARI measures the similarity between two different partitions and ranges from −1 to 1. A larger value of ARI means a higher degree of similarity between two partitions. ARI takes a value of 1 when the two partitions are equal up to a permutation. McFadden’s pseudo-R² and DB-Index were used to measure the performance of the feature extraction methods. McFadden’s pseudo-R² [70] is an evaluation metric used in the context of statistical models, particularly in the assessment of the goodness-of-fit for logistic regression models. It provides a measure of how well the model explains the variability in the data. A higher pseudo R² value indicates that the model explains a larger proportion of the variance, suggesting a better fit. DB-index is a metric used to evaluate the quality of clustering algorithms by measuring the compactness and separation between clusters. It is based on the ratio of within-cluster scatter to between-cluster separation, providing an assessment of how well the clusters are formed.

Lower DB-index values indicate better feature extraction performance, as they suggest more compact and well-separated group in the feature reduced space. For intuitive interpretation, the DB-Index is presented as 1 – DB-Index, converting the metric so that higher values correspond directly to better clustering performance.

### Data sets

#### Mouse preoptic hypothalamus 3D MERFISH dataset

Spatial transcriptomic data for the mouse preoptic hypothalamus obtained using MERFISH technology was retrieved from the BASS GitHub repository: https://zhengli09.github.io/BASS-Analysis/MERFISH.html. The original data are accessible from the study by Moffitt et al. [13]. Li and Zhou [25] retrieved the data from five tissue sections of the preoptic hypothalamus from a single mouse, corresponding to Bregma coordinates −0.04, −0.09, −0.14, −0.19, and −0.24 mm. Briefly, the raw expression count matrix was analyzed and quality control was performed QC on each sample, filtering out genes with non-zero expression levels in fewer than 20 spots, and spots with non-zero expression levels in fewer than 20 genes. The annotated spatial domains for each section were manually defined based on known anatomical landmarks in the original study. In the analysis, these annotations served as the ground truth to evaluate the clustering and data integration performance of different methods.

Subsequently, the top 100 spatially variable genes were selected using SPARK-X independently for each section. PCA was performed to reduce the dimensionality of the data (with and without passing-message procedure) to 20 principal components.

#### DLPFC 10X Visium dataset

The ST data from the human DLPFC, acquired via the 10X Visium platform, were obtained as a SpatialExperiment object using the R package spatialLIBD. Prior to analysis, spots were filtered based on quality control criteria (library size <500, number of detected genes <250, mitochondrial gene percentage >30, and cell count >12). Genes were retained if they exhibited at least 2 counts in at least 0.5% of spatial locations, and all mitochondrial genes were excluded. This filtering process resulted in an average of 2582 genes across a total of 46318 spatial locations after removal of non-labeled spots. The expression matrix was normalized using the logNormCounts function. Subsequently, the top 2000 spatially variable genes were selected using SPARK-X independently for each section. PCA was performed to reduce the dimensionality of the data to 50 principal components.

#### 10X Visium Human prostate tissues

Data can be downloaded from the 10x website (“Normal Prostate”, “Acinar Cell Carcinoma”, “Adjacent Normal Section”, and “Adenocarcinoma”). A quality control step is performed to filter out low-quality spots based on the number of detected transcripts and mitochondrial genes. Genes with low expression in the spots were filtered out, retaining only those with sufficient expression levels for analysis. The counts were normalized using library size factors, and a logarithmic transformation was applied to prepare the data for downstream analyses.

Subsequently, the top 2000 spatially variable genes were selected using SPARK-X independently for each section. PCA was performed to reduce the dimensionality of the data to 50 principal components. Harmony algorithm [45] was applied to correct the batch effects related to sample identity.

#### Human Colorectal Cancer Visium HD dataset

Data can be downloaded from the 10x website. The data undergoes preprocessing and subsequent analytical steps using the Seurat pipeline. Initially, the data is loaded into R using the Load10X_Spatial function. A quality control step is performed to filter out low-quality spots based on the number of detected transcripts. This step ensures that only high-quality ST data is retained for further analysis. Following quality control, the preprocessing proceeds with normalization and identification of variable features using Seurat functions. Subsequently, the top 2000 spatially variable genes were selected using SPARK-X independently for each section. PCA was performed to reduce the dimensionality of the data to 50 principal components.

### Feature extraction and clustering

#### Louvain and Leiden

Louvain [71] and Leiden [72] are nonspatial clustering algorithms that allocate each spot to a distinct community and optimize modularity by iteratively merging and splitting communities until the desired clustering results are achieved. Leiden can also merge similar communities to enhance clustering results further. Both these methods have been adopted into the Seurat pipeline.

#### Walktrap

Walktrap [73] is a nonspatial clustering algorithm based on the observation that short random walks are more likely to remain within the same community rather than crossing into different communities. The walktrap algorithm is particularly effective for detecting communities with strong local structures and provides a hierarchical decomposition of the network, enabling multi-resolution analysis of the community structure. The algorithm has been adopted in the Giotto pipeline.

#### BayesSpace

BayesSpace [74] is a full Bayesian statistical method incorporating a low-dimensional representation of the gene expression matrix to model spatial clustering, encouraging neighboring points to belong to the same cluster through a prior spatial prior. This method requires spatial data in spot locations, so it is unsuitable for data that do not use ‘spot’.

#### BASS

BASS [25] is a spatial clustering method that facilitates multiscale and multisample analysis of ST data. It utilizes a Bayesian hierarchical modeling framework for clustering analysis.

#### BANKSY

BANKSY [30] is a spatially aware clustering algorithm designed to leverage both gene expression and spatial information to refine biological insights. It incorporates spatial constraints in clustering to improve accuracy in detecting functional tissue regions.

#### PRECAST

PRECAST [29] is a spatial clustering framework that integrates multimodal ST data. It employs regularized factor models to identify distinct spatial domains, providing a robust and scalable solution for analyzing large-scale ST datasets.

### Statistical analysis

Differences in numerical variables between two groups were assessed using the Wilcoxon rank-sum test. Correlation between variables was evaluated using Pearson’s and Spearman’s correlation coefficients. To identify complex, non-linear associations without assuming a specific functional relationship, the MIC [75]was calculated. Statistical significance was defined as a p-value < 0.05. Where applicable, false discovery rate (FDR) correction was applied to account for multiple testing.

### Gene set enrichment analysis (GSEA)

GSEA for spatial transcriptomic data was performed using, respectively, the variance-adjusted Mahalanobis (VAM) method [76] on multiple gene sets obtained from the hallmarks gene set collection [77]. VAM method implemented in the R VAM package computes distance statistics and one-sided p-values for all cells. The p-values were computed using gamma distribution.

### Image processing

Histopathological microscopy images, including those of prostate tissues and colorectal carcinoma, were downloaded from the 10x Genomics dataset repository (https://www.10xgenomics.com/datasets). These images were subsequently converted into pyramidal open microscopy environment (OME) tagged image file (TIF) format using ImageJ (version 2.16.0) and its plugin Kheops [78] to facilitate efficient visualization and analysis. The histopathology microscopy image of the DLPFC dataset were obtained from the publicly available repository hosted by the Lieber Institute (https://github.com/LieberInstitute/HumanPilot). Spatially aligned image tiles corresponding to individual Visium spots were generated utilizing a Python script leveraging the SpatialData framework [55] and stored as 128×128 TIFF images.

### Digital histological analysis

A CNN was designed for image classification and structured using Keras [79] in R. TIFF images were loaded as a four-dimensional tensor suitable for convolutional input. The model included two convolutional blocks with ReLU activation (16 or 32 filters), followed by 2×2 max pooling. Instead of flattening, global average pooling condensed feature maps, and a final dense layer with softmax classified the tiles. The model was trained for 20 epochs using the Adam optimizer and sparse categorical crossentropy loss. Performance was assessed on the test set via accuracy, loss, and a confusion matrix comparing predictions to true labels.

The model was embedded in an MC cross-validation procedure where 75% of the tiles were included in the balanced training set (2566 tiles for sublayer 1 and 2566 tiles for sublayers 3 and 5) and the rest 25% were included in the test set. The procedure was repeated 100 times to estimate an average (and standard deviation) of the prediction accuracy.

## Data availability

All spatial transcriptomics datasets used in this study are publicly available. The 3D MERFISH mouse hypothalamus dataset was obtained from the BASS GitHub repository (https://zhengli09.github.io/BASS-Analysis/MERFISH.html) and originally published by Moffitt *et al.* [13] The DLPFC dataset was accessed through the spatialLIBD R package (http://spatial.libd.org/spatialLIBD/). The 10x Visium human prostate and colorectal cancer (Visium HD) datasets were downloaded from the 10x Genomics website (https://www.10xgenomics.com/resources/datasets). Processed data matrices, intermediate results, and clustering outputs generated during the current study are available upon request or through the KODAMA analysis repository at https://tkcaccia.github.io/KODAMA-Analysis.

## Code availability

The KODAMA software code is publicly available at https://github.com/tkcaccia/KODAMA and https://github.com/tkcaccia/KODAMAextra. The source code is released under the GNU General Public License version 3 (GPL >=3). All analysis codes for reproducing the results of the present study are publicly available at https://tkcaccia.github.io/KODAMA-Analysis/

## Acknowledgements

The authors thank the University of Cape Town’s ICTS High Performance Computing team for providing a high-performance computing facility for this study (https://ucthpc.uct.ac.za/).

## Supplementary information

**Supplementary Table 1:** Differentially expressed genes among DLPFC layer 3 subclusters. This table lists genes significantly overexpressed in clusters 2, 3, and 4 identified within cortical layer 3 by KODAMA in the DLPFC dataset. Median expression values and 95% confidence intervals are provided for each cluster, along with p-values and false discovery rates (FDRs) for pairwise comparisons.

| Gene overexpressed in Cluster 2 | Cluster 2 | Cluster 3 | Cluster 4 | Cluster 2 vs. Cluster 3 |  | Cluster 2 vs. Cluster 4 |  |
| --- | --- | --- | --- | --- | --- | --- | --- |
|  |  |  |  | p-value | FDR | p-value | FDR |
| MALAT1, median [95%CI] | 3.47 [0 7.60] | 2.75 [0 6.62] | 2.33 [0 5.30] | 1.53E-41 | 1.02E-38 | 7.83E-74 | 5.22E-71 |
| CAMK2N1, median [95%CI] | 6.21 [0 8.70] | 5.57 [0 7.85] | 5.03 [0 7.47] | 1.58E-49 | 1.58E-46 | 3.93E-106 | 3.93E-103 |
| HPCAL1, median [95%CI] | 2.77 [0 6.81] | 0 [0 5.48] | 0 [0 4.37] | 8.54E-75 | 1.71E-71 | 1.02E-107 | 2.03E-104 |
| PPP3CA, median [95%CI] | 3.93 [0 7.58] | 3.42 [0 6.63] | 3.12 [0 6.30] | 1.37E-17 | 3.44E-15 | 1.07E-22 | 1.79E-20 |
| ARPP19, median [95%CI] | 4.69 [0 7.68] | 4.21 [0 7.01] | 3.75 [0 6.33] | 6.25E-22 | 3.13E-19 | 3.30E-53 | 1.10E-50 |
| EEF1A1, median [95%CI] | 6.24 [2.14 8.6] | 5.85 [2.40 8.07] | 5.71 [2.65 7.84] | 1.84E-19 | 7.36E-17 | 9.57E-33 | 2.13E-30 |
| PEA15, median [95%CI] | 3.47 [0 7.21] | 2.92 [0 6.43] | 2.68 [0 5.99] | 2.06E-17 | 4.58E-15 | 4.47E-21 | 6.38E-19 |
| RPL4, median [95%CI] | 4.97 [0 8.36] | 4.49 [0 7.74] | 4.35 [0 7.40] | 1.27E-15 | 2.30E-13 | 4.32E-17 | 4.55E-15 |
| NPTXR, median [95%CI] | 2.88 [0 6.94] | 2.24 [0 6.08] | 2.23 [0 5.77] | 5.37E-11 | 8.26E-09 | 1.56E-13 | 1.20E-11 |
| ENC1, median [95%CI] | 5.65 [0 8.3] | 5.34 [0 7.88] | 4.67 [0 7.21] | 6.12E-11 | 8.75E-09 | 1.58E-68 | 7.92E-66 |

| Gene overexpressed in Cluster 3 | Cluster 2 | Cluster 3 | Cluster 4 | Cluster 3 vs. Cluster 2 |  | Cluster 3 vs. Cluster 4 |  |
| --- | --- | --- | --- | --- | --- | --- | --- |
|  |  |  |  | p-value | FDR | p-value | FDR |
| YWHAH, median [95%CI] | 5.77 [0 8.15] | 5.97 [0.54 8.06] | 5.72 [2.27 7.71] | 1.18E-05 | 7.00E-05 | 1.55E-07 | 2.58E-05 |
| CCK, median [95%CI] | 5.08 [0 7.06] | 5.22 [2.26 6.87] | 4.98 [2.16 6.6] | 1.38E-03 | 4.34E-03 | 2.40E-07 | 3.43E-05 |
| PAK1, median [95%CI] | 2.07 [0 6.21] | 2.53 [0 5.95] | 2.28 [0 5.40] | 2.29E-05 | 1.25E-04 | 4.11E-03 | 1.65E-01 |
| RGS4, median [95%CI] | 3.11 [0 6.26] | 3.23 [0 6.09] | 3.00 [0 5.74] | 1.72E-03 | 5.22E-03 | 6.10E-03 | 2.20E-01 |
| MT2A, median [95%CI] | 2.93 [0 6.69] | 3.00 [0 6.62] | 2.69 [0 6.11] | 1.52E-02 | 3.33E-02 | 1.87E-03 | 8.74E-02 |
| CALM2, median [95%CI] | 6.64 [2.77 8.74] | 6.81 [3.23 8.66] | 6.71 [3.71 8.51] | 8.93E-08 | 9.57E-07 | 2.08E-02 | 5.33E-01 |
| APLP2, median [95%CI] | 2.82 [0 7.02] | 3.18 [0 6.87] | 2.88 [0 6.42] | 4.54E-05 | 2.24E-04 | 2.69E-02 | 6.56E-01 |
| VSNL1, median [95%CI] | 6.21 [0 8.49] | 6.29 [2.68 8.36] | 6.15 [2.53 8.16] | 2.86E-02 | 5.67E-02 | 7.62E-03 | 2.51E-01 |
| SNX3, median [95%CI] | 3.18 [0 7.24] | 3.22 [0 6.72] | 2.91 [0 6.38] | 4.85E-02 | 8.87E-02 | 3.21E-04 | 2.07E-02 |
| ITM2C, median [95%CI] | 3.97 [0 7.33] | 4.11 [0 7.35] | 4.02 [0 6.91] | 6.14E-03 | 1.55E-02 | 6.02E-02 | 1.00E+00 |

| Gene overexpressed in Cluster 4 | Cluster 2 | Cluster 3 | Cluster 4 | Cluster 4 vs. Cluster 2 |  | Cluster 4 vs. Cluster 3 |  |
| --- | --- | --- | --- | --- | --- | --- | --- |
|  |  |  |  | p-value | FDR | p-value | FDR |
| MBP, median [95%CI] | 4.04 [0 7.60] | 4.73 [0 7.96] | 5.35 [0 8.19] | 2.11E-89 | 7.03E-87 | 2.08E-21 | 1.38E-18 |
| TUBA1B, median [95%CI] | 5.98 [0 8.27] | 6.79 [3.65 8.63] | 7.16 [4.62 8.74] | 2.32E-174 | 4.64E-171 | 4.30E-25 | 4.30E-22 |
| SCN1B, median [95%CI] | 0 [0 6.61] | 2.75 [0 6.86] | 3.40 [0 7.1] | 6.42E-98 | 3.21E-95 | 3.23E-18 | 1.62E-15 |
| NEFL, median [95%CI] | 3.95 [0 7.34] | 5.29 [0 8.13] | 5.80 [0 8.28] | 9.52E-153 | 9.52E-150 | 1.67E-13 | 5.57E-11 |
| QDPR, median [95%CI] | 0 [0 6.81] | 1.87 [0 7.11] | 3.01 [0 7.02] | 2.05E-50 | 2.93E-48 | 5.33E-13 | 1.52E-10 |
| CNP, median [95%CI] | 0 [0 6.10] | 0 [0 5.85] | 2.43 [0 6.09] | 1.49E-25 | 5.74E-24 | 1.83E-12 | 4.08E-10 |
| LGALS1, median [95%CI] | 2.43 [0 6.56] | 3.33 [0 6.87] | 3.75 [0 7.02] | 5.43E-79 | 1.09E-76 | 5.41E-12 | 9.85E-10 |
| KRT19, median [95%CI] | 0 [0 6.18] | 0 [0 6.25] | 2.09 [0 6.46] | 3.02E-31 | 1.72E-29 | 9.91E-11 | 1.52E-08 |
| PLP1, median [95%CI] | 2.53 [0 6.80] | 3.05 [0 7.04] | 3.39 [0 7.30] | 8.88E-48 | 1.04E-45 | 2.38E-10 | 3.17E-08 |
| TMSB10, median [95%CI] | 6.67 [3.40 8.72] | 7.06 [4.48 8.87] | 7.26 [5.16 8.85] | 5.21E-65 | 9.47E-63 | 2.98E-10 | 3.72E-08 |

**Supplementary Table 2:**
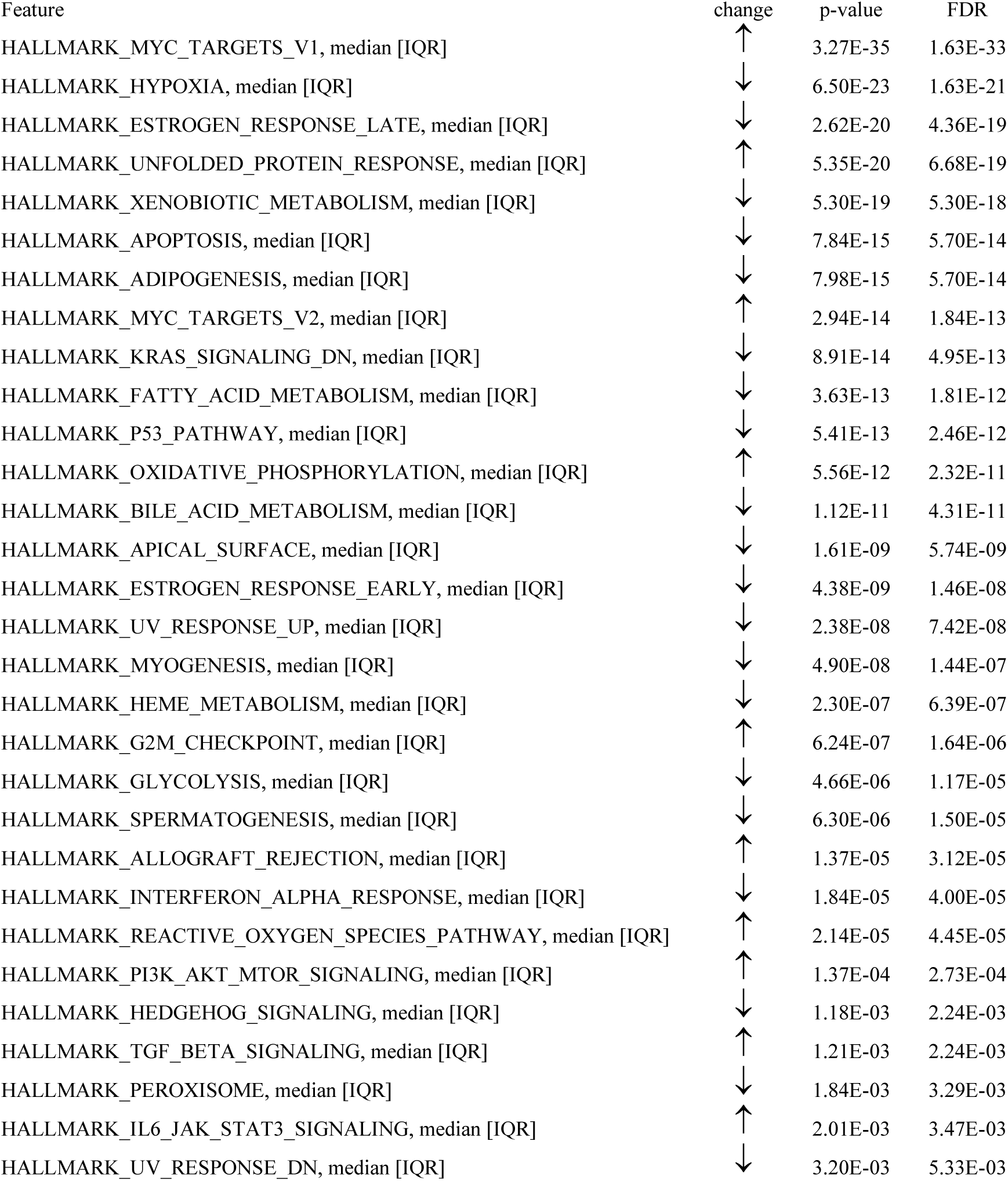
Differentially enriched gene sets between normal- and tumor-like phenotypes in prostate tissue, defined as “normal”. GSEA comparing regions histologically labeled as normal but exhibiting tumor-like gene expression patterns, identified by KODAMA. Gene sets from the MSigDB Hallmark collection are listed with direction of change, p-values, and FDRs. Notable changes include upregulation of MYC targets and downregulation of hypoxia and estrogen response pathways.

| Feature | change | p-value | FDR |
| --- | --- | --- | --- |
| HALLMARK_MYC_TARGETS_V1, median [IQR] | ↑ | 3.27E-35 | 1.63E-33 |
| HALLMARK_HYPOXIA, median [IQR] | ↓ | 6.50E-23 | 1.63E-21 |
| HALLMARK_ESTROGEN_RESPONSE_LATE, median [IQR] | ↓ | 2.62E-20 | 4.36E-19 |
| HALLMARK_UNFOLDED_PROTEIN_RESPONSE, median [IQR] | ↑ | 5.35E-20 | 6.68E-19 |
| HALLMARK_XENOBIOTIC_METABOLISM, median [IQR] | ↓ | 5.30E-19 | 5.30E-18 |
| HALLMARK_APOPTOSIS, median [IQR] | ↓ | 7.84E-15 | 5.70E-14 |
| HALLMARK_ADIPOGENESIS, median [IQR] | ↓ | 7.98E-15 | 5.70E-14 |
| HALLMARK_MYC_TARGETS_V2, median [IQR] | ↑ | 2.94E-14 | 1.84E-13 |
| HALLMARK_KRAS_SIGNALING_DN, median [IQR] | ↓ | 8.91E-14 | 4.95E-13 |
| HALLMARK_FATTY_ACID_METABOLISM, median [IQR] | ↓ | 3.63E-13 | 1.81E-12 |
| HALLMARK_P53_PATHWAY, median [IQR] | ↓ | 5.41E-13 | 2.46E-12 |
| HALLMARK_OXIDATIVE_PHOSPHORYLATION, median [IQR] | ↑ | 5.56E-12 | 2.32E-11 |
| HALLMARK_BILE_ACID_METABOLISM, median [IQR] | ↓ | 1.12E-11 | 4.31E-11 |
| HALLMARK_APICAL_SURFACE, median [IQR] | ↓ | 1.61E-09 | 5.74E-09 |
| HALLMARK_ESTROGEN_RESPONSE_EARLY, median [IQR] | ↓ | 4.38E-09 | 1.46E-08 |
| HALLMARK_UV_RESPONSE_UP, median [IQR] | ↓ | 2.38E-08 | 7.42E-08 |
| HALLMARK_MYOGENESIS, median [IQR] | ↓ | 4.90E-08 | 1.44E-07 |
| HALLMARK_HEME_METABOLISM, median [IQR] | ↓ | 2.30E-07 | 6.39E-07 |
| HALLMARK_G2M_CHECKPOINT, median [IQR] | ↑ | 6.24E-07 | 1.64E-06 |
| HALLMARK_GLYCOLYSIS, median [IQR] | ↓ | 4.66E-06 | 1.17E-05 |
| HALLMARK_SPERMATOGENESIS, median [IQR] | ↓ | 6.30E-06 | 1.50E-05 |
| HALLMARK_ALLOGRAFT_REJECTION, median [IQR] | ↑ | 1.37E-05 | 3.12E-05 |
| HALLMARK_INTERFERON_ALPHA_RESPONSE, median [IQR] | ↓ | 1.84E-05 | 4.00E-05 |
| HALLMARK_REACTIVE_OXYGEN_SPECIES_PATHWAY, median [IQR] | ↑ | 2.14E-05 | 4.45E-05 |
| HALLMARK_PI3K_AKT_MTOR_SIGNALING, median [IQR] | ↑ | 1.37E-04 | 2.73E-04 |
| HALLMARK_HEDGEHOG_SIGNALING, median [IQR] | ↓ | 1.18E-03 | 2.24E-03 |
| HALLMARK_TGF_BETA_SIGNALING, median [IQR] | ↑ | 1.21E-03 | 2.24E-03 |
| HALLMARK_PEROXISOME, median [IQR] | ↓ | 1.84E-03 | 3.29E-03 |
| HALLMARK_IL6_JAK_STAT3_SIGNALING, median [IQR] | ↑ | 2.01E-03 | 3.47E-03 |
| HALLMARK_UV_RESPONSE_DN, median [IQR] | ↓ | 3.20E-03 | 5.33E-03 |

**Supplementary Table 3:** Genes associated with the spatial trajectory in prostate cancer. This table reports genes correlated with a manually defined trajectory tracing tumor development from core to periphery in the prostate adenocarcinoma sample. Genes such as SAA2 and NNMT are highlighted for their potential roles in early tumorigenesis and immune modulation.

| **Feature** | **rho** | **p-value** | **FDR** |
| --- | --- | --- | --- |
| LCN2 | -0.88 | 6.92E-21 | 6.94E-18 |
| SOD2 | -0.81 | 3.83E-15 | 1.92E-12 |
| CEBPD | -0.81 | 5.75E-15 | 1.92E-12 |
| CXCL3 | -0.77 | 7.34E-13 | 1.67E-10 |
| ID1 | -0.77 | 8.35E-13 | 1.67E-10 |
| IL32 | -0.75 | 4.17E-12 | 6.97E-10 |
| PI3 | -0.74 | 1.34E-11 | 1.91E-09 |
| CCL20 | -0.74 | 1.73E-11 | 2.17E-09 |
| CXCL1 | -0.74 | 2.16E-11 | 2.40E-09 |
| TRIM31 | -0.73 | 2.89E-11 | 2.90E-09 |
| NCOA7 | -0.73 | 3.50E-11 | 3.19E-09 |
| CXCL2 | -0.73 | 4.02E-11 | 3.36E-09 |
| HK2 | -0.71 | 1.76E-10 | 1.28E-08 |
| GPX2 | -0.71 | 1.78E-10 | 1.28E-08 |
| NOS2 | -0.7 | 4.64E-10 | 3.06E-08 |
| TNFRSF21 | -0.7 | 4.89E-10 | 3.06E-08 |
| MUC1 | -0.69 | 8.34E-10 | 4.92E-08 |
| TRIB1 | -0.69 | 1.16E-09 | 6.49E-08 |
| CFB | -0.68 | 1.87E-09 | 9.85E-08 |
| S100P | -0.68 | 1.99E-09 | 9.97E-08 |
| SPINK1 | -0.67 | 4.15E-09 | 1.98E-07 |
| CKB | -0.67 | 6.05E-09 | 2.76E-07 |
| CEACAM1 | -0.66 | 7.66E-09 | 3.34E-07 |
| ERO1A | -0.66 | 9.16E-09 | 3.83E-07 |
| FOS | -0.65 | 2.38E-08 | 9.50E-07 |
| AREG | -0.65 | 2.47E-08 | 9.50E-07 |
| TDP2 | -0.65 | 2.56E-08 | 9.50E-07 |
| WARS | -0.64 | 3.70E-08 | 1.33E-06 |
| BSPRY | -0.64 | 4.82E-08 | 1.67E-06 |
| SELENBP1 | -0.63 | 6.07E-08 | 2.03E-06 |

**Supplementary Table 4:**
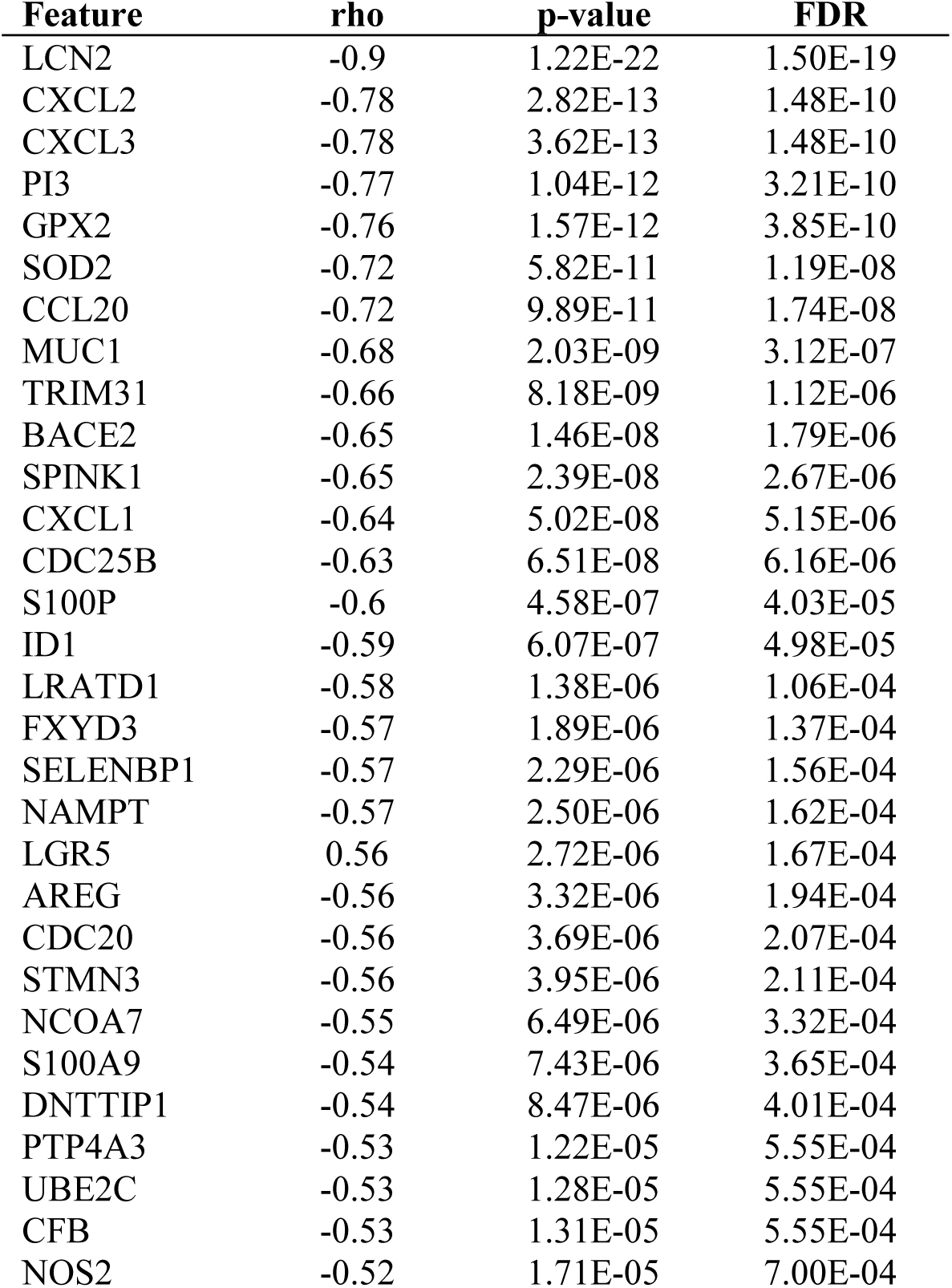
Genes associated with the KODAMA-inferred gradient in colorectal carcinoma. A ranked list of genes showing expression gradients along a trajectory identified by KODAMA in the VisiumHD colorectal tumor dataset. Genes are ordered by strength of association with spatial progression, with CXCL2 among the top candidates involved in tumor invasion and microenvironment remodeling.

**Supplementary Table 5:** Genes associated with distance from invasive carcinoma. Genes identified as highly correlated with proximity to invasive carcinoma regions in colorectal tissue, based on the MIC value. These include IGFBP5, MGP, and HTRA3, which are implicated in extracellular matrix remodeling and tumor progression.

| **Feature** | **MIC** | **p-value** | **FDR** |
| --- | --- | --- | --- |
| IGFBP5 | 1.00 | 3.90E-239 | 3.92E-237 |
| HTRA3 | 1.00 | 9.10E-261 | 1.83E-258 |
| MGP | 0.99 | 7.32E-191 | 3.68E-189 |
| TIMP3 | 0.92 | 1.88E-171 | 6.31E-170 |
| GREM1 | 0.91 | 4.30E-206 | 2.88E-204 |
| IGFBP3 | 0.89 | 4.32E-176 | 1.74E-174 |
| SFRP4 | 0.86 | 1.84E-170 | 5.29E-169 |
| CXCL14 | 0.84 | 9.84E-148 | 1.80E-146 |
| COL1A1 | 0.83 | 7.49E-152 | 1.67E-150 |
| MMP11 | 0.8 | 1.07E-141 | 1.66E-140 |
| SPARC | 0.76 | 1.99E-128 | 2.50E-127 |
| AEBP1 | 0.76 | 6.24E-99 | 5.23E-98 |
| CCDC80 | 0.75 | 2.47E-129 | 3.31E-128 |
| SFRP2 | 0.75 | 3.47E-90 | 2.49E-89 |
| ISLR | 0.75 | 3.13E-161 | 7.88E-160 |
| COL1A2 | 0.74 | 7.60E-102 | 7.27E-101 |
| DCN | 0.74 | 5.30E-98 | 4.26E-97 |
| COL3A1 | 0.73 | 6.96E-132 | 9.99E-131 |
| COL14A1 | 0.73 | 2.01E-100 | 1.83E-99 |
| A2M | 0.73 | 2.73E-99 | 2.38E-98 |
| COL11A1 | 0.72 | 1.90E-143 | 3.18E-142 |
| LUM | 0.71 | 1.27E-150 | 2.55E-149 |
| COMP | 0.71 | 5.95E-109 | 6.64E-108 |
| IGKC | 0.70 | 4.51E-117 | 5.34E-116 |
| ELN | 0.70 | 2.16E-85 | 1.36E-84 |
| ANGPTL2 | 0.70 | 1.52E-92 | 1.18E-91 |
| ITGBL1 | 0.69 | 1.58E-103 | 1.67E-102 |
| IGHG1 | 0.68 | 1.75E-88 | 1.21E-87 |
| MMP2 | 0.68 | 1.28E-87 | 8.59E-87 |
| FN1 | 0.67 | 2.31E-102 | 2.32E-101 |

**Supplementary Figure 1:**
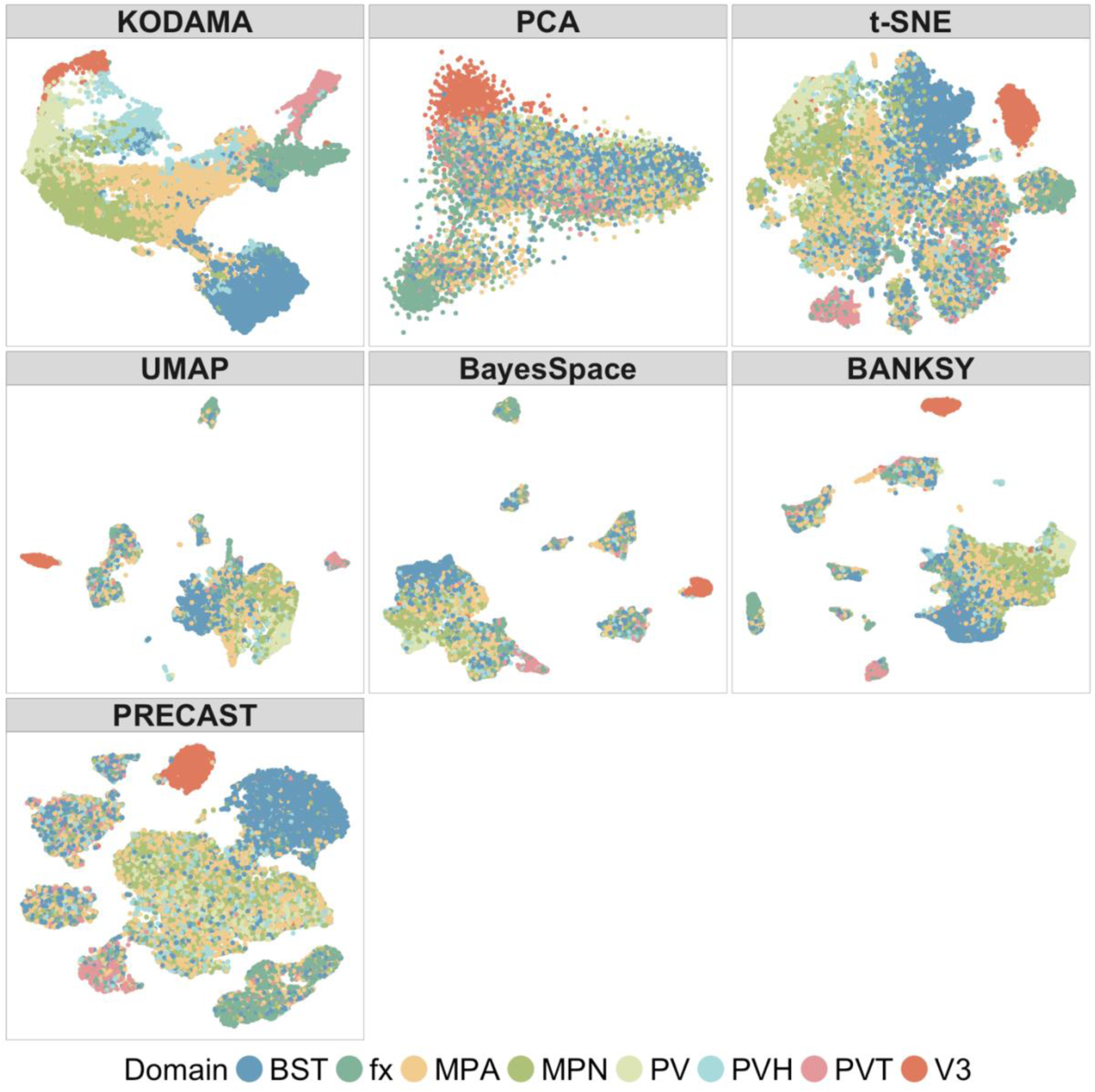
Comparison of feature extraction methods across tissue sections. Visual representation of dimensionality reduction results (*i.e.*, KODAMA, PCA, *t*-SNE, UMAP, BayesSpace, BANKSY, and PRECAST) applied to the tissue sections of the MERFISH dataset, highlighting differences in spatial coherence and structure preservation.

**Supplementary Figure 2:**
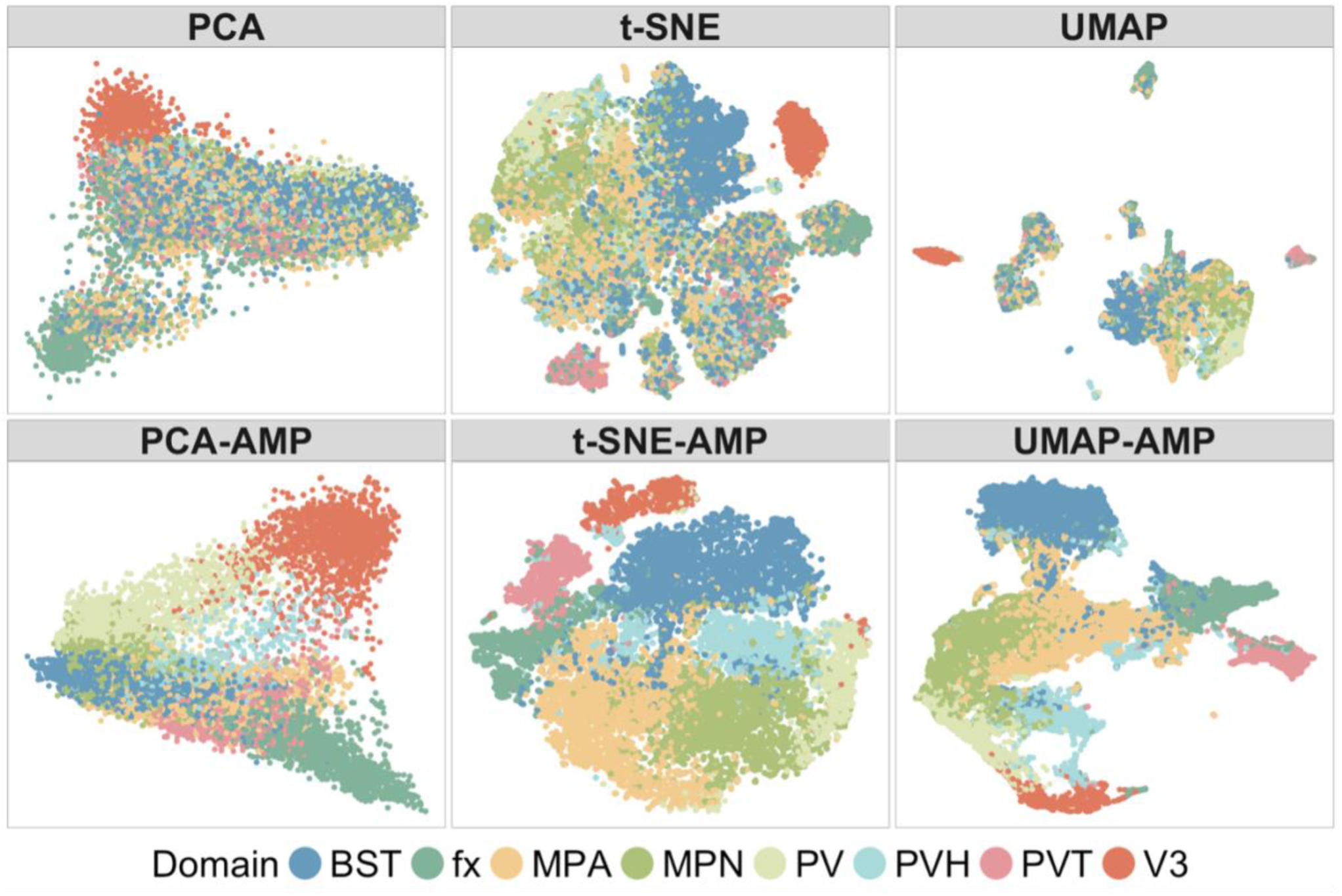
Impact of AMP preprocessing on non-spatial methods. Side-by-side comparison of PCA, UMAP, and *t*-SNE with and without AMP, demonstrating how AMP enhances the separation of biological domains in spatial transcriptomics data.

**Supplementary Figure 3:**
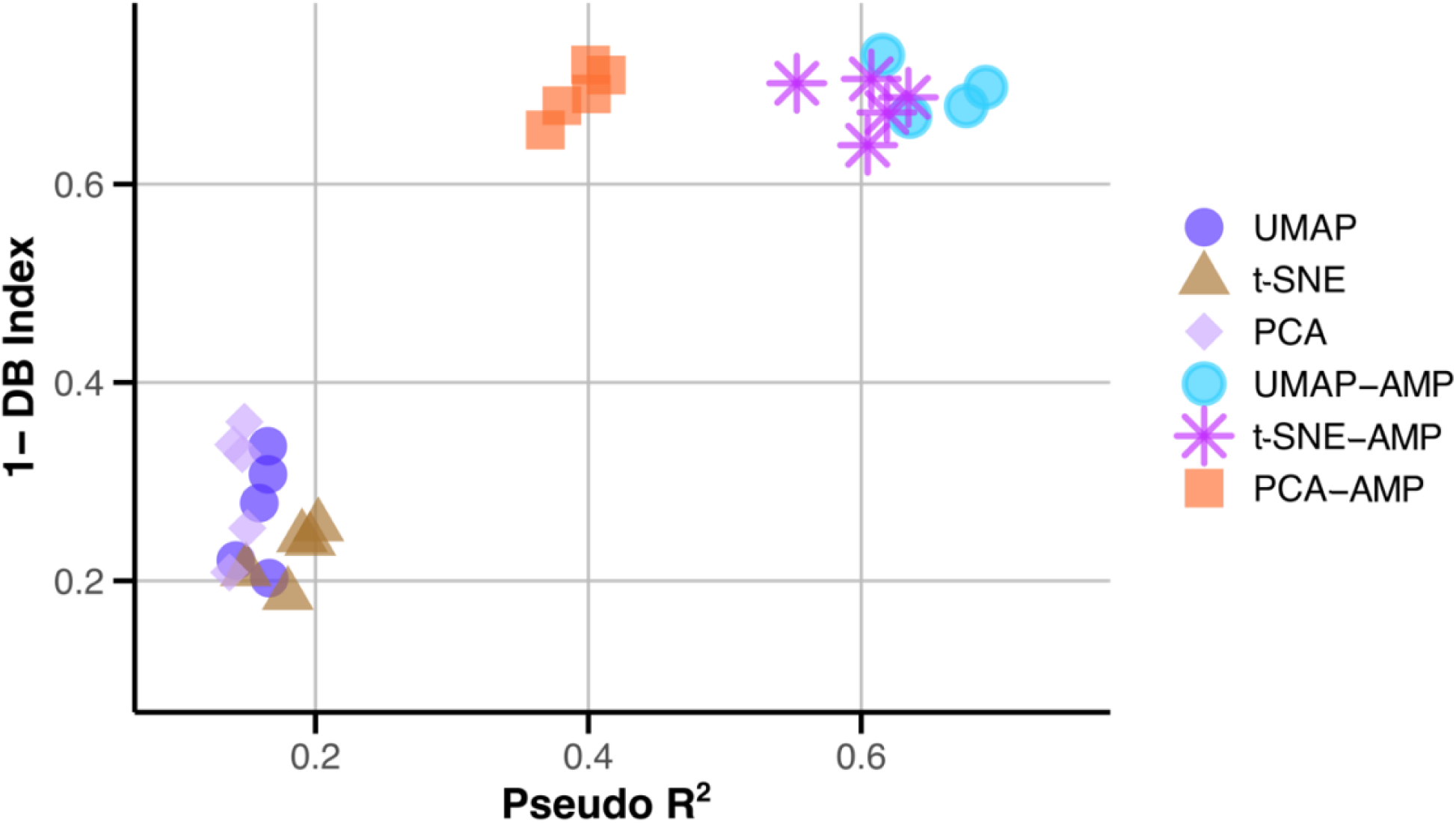
Quantitative assessment of feature extraction methods. Boxplots of inverse Davies–Bouldin Index (1–DB Index) and McFadden’s pseudo-R² scores across multiple feature extraction methods, with and without AMP preprocessing.

**Supplementary Figure 4:**
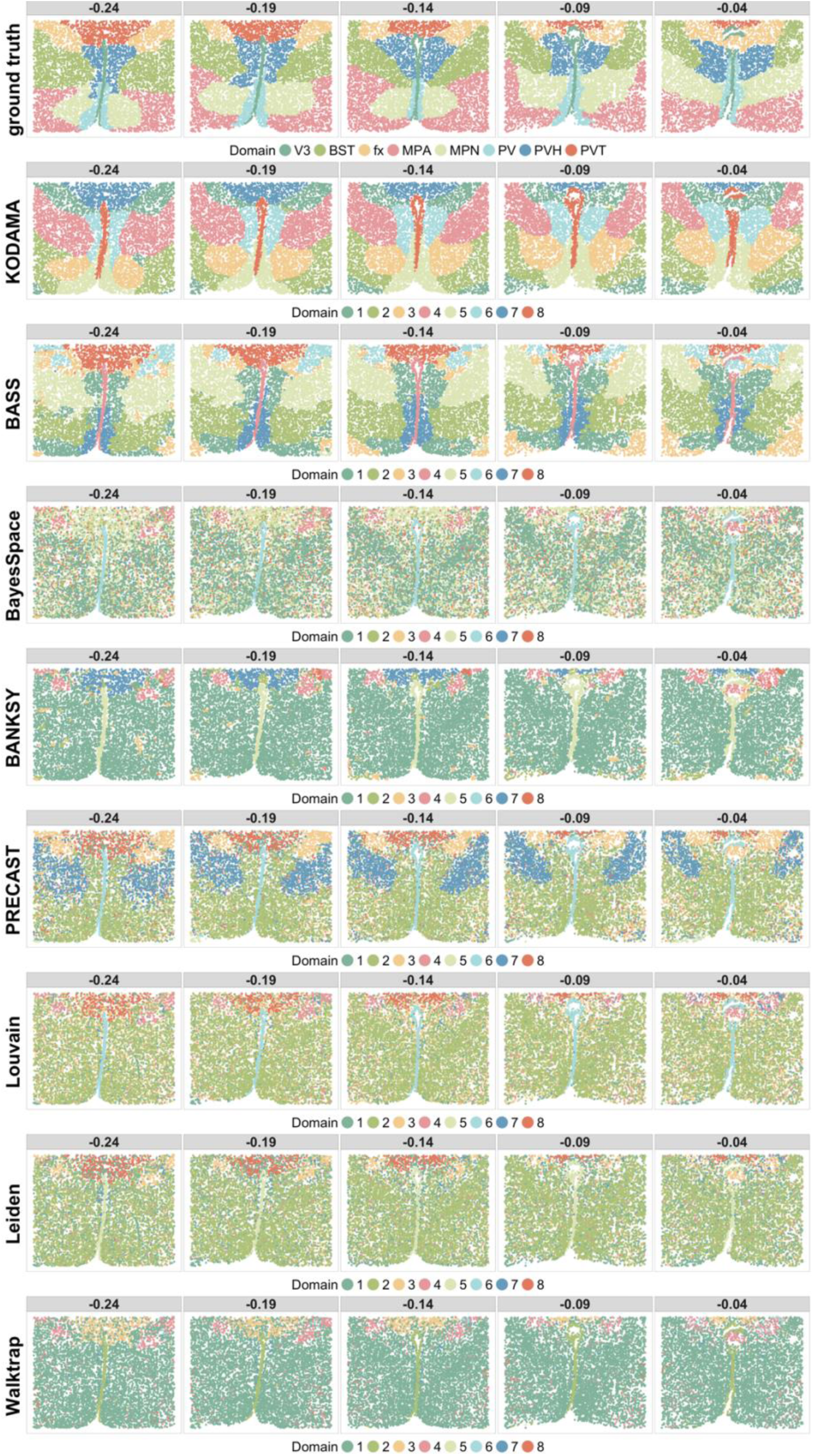
Benchmarking KODAMA against other spatial and non-spatial clustering methods on MERFISH data. Comparison of KODAMA with spatial (*i.e.*, BASS, BayesSpace, BANKSY, PRECAST) and non-spatial (*i.e.*, Louvain, Leiden, Walktrap) clustering algorithms, showing the spatial domain recovery and structural accuracy.

**Supplementary Figure 5:**
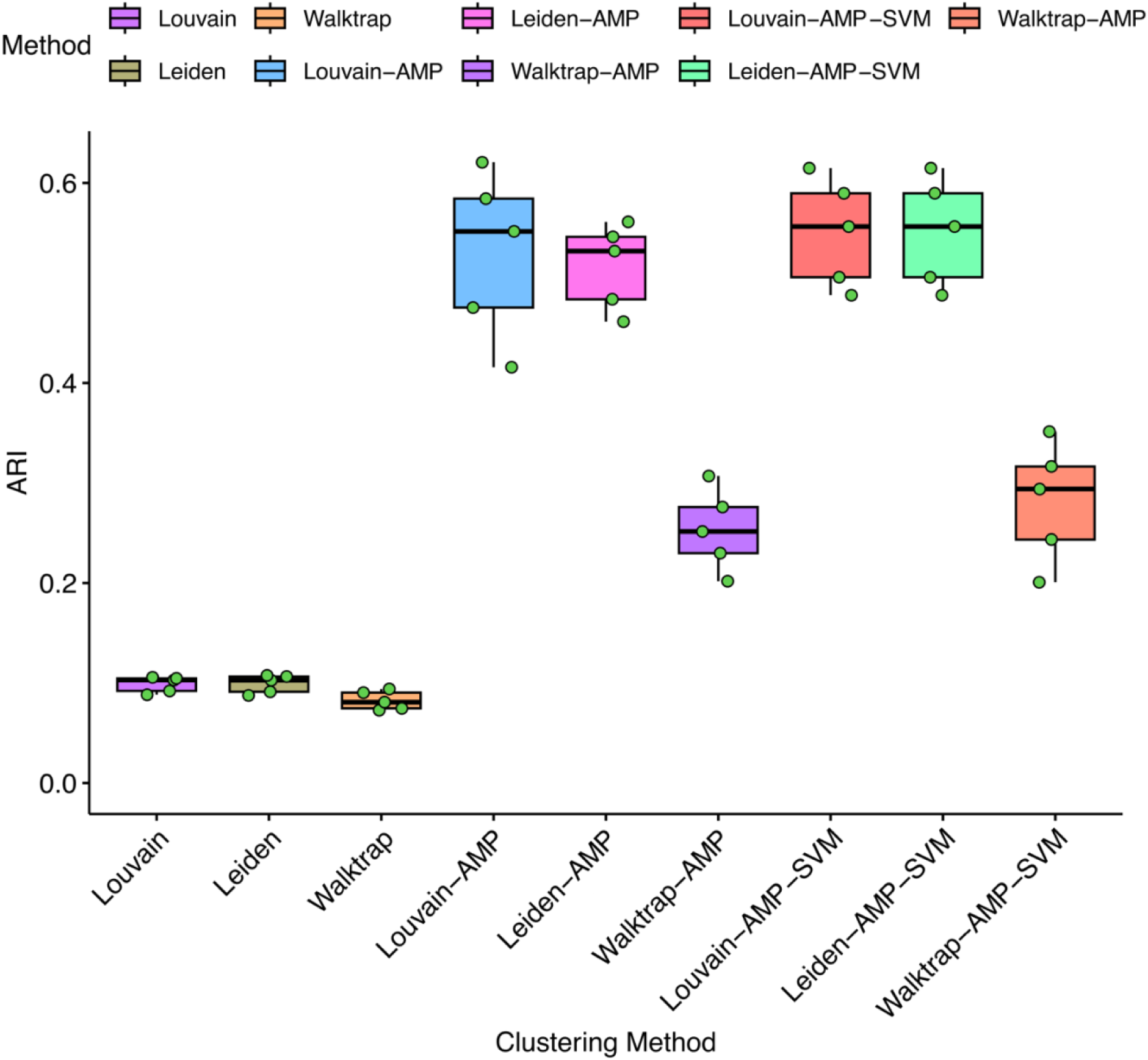
Enhancement of non-spatial clustering via message-passing and SVM refinement. Improved tissue segment identification in non-spatial methods following the integration of AMP and SVM-based label refinement, applied to MERFISH datasets.

**Supplementary Figure 6:**
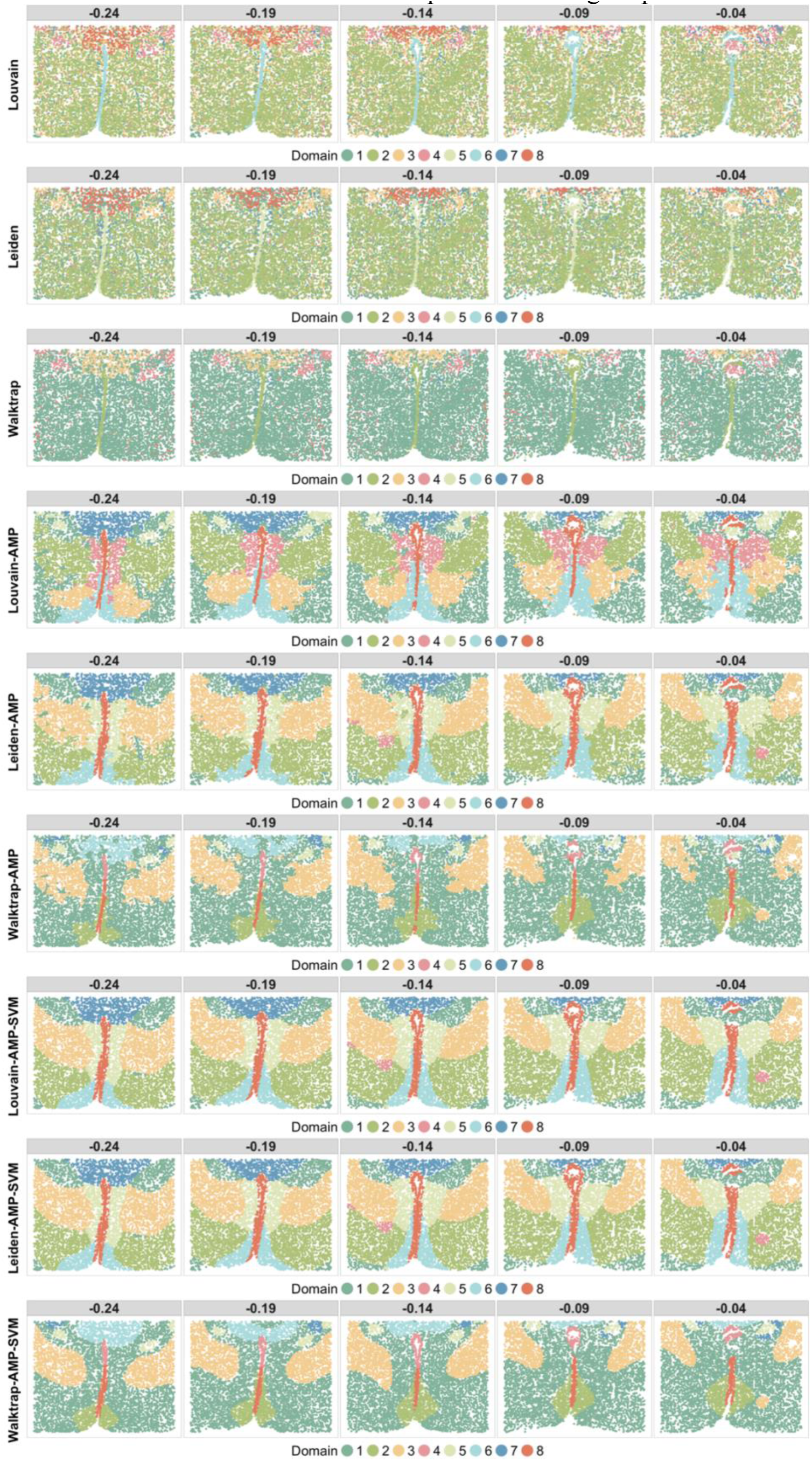
Replication of clustering improvement using AMP and SVM. Confirmation that message-passing and SVM post-processing consistently improve spatial domain detection across different non-spatial clustering outputs.

**Supplementary Figure 7:**
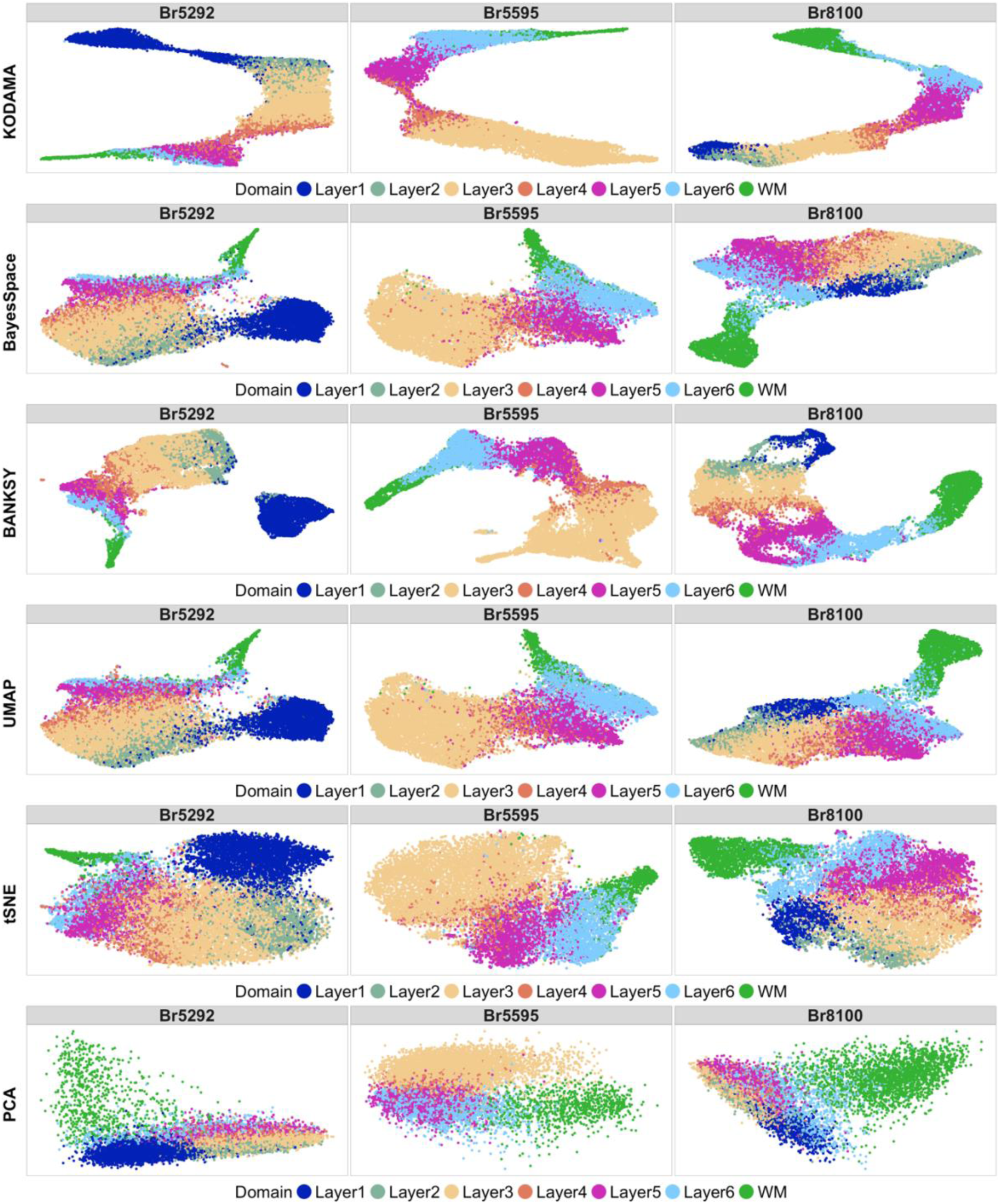
Feature extraction outputs for DLPFC tissue. Visualization of the dimensionality reduction results from various methods applied to 12 human DLPFC tissue sections, highlighting differences in spatial layer separation.

**Supplementary Figure 8:**
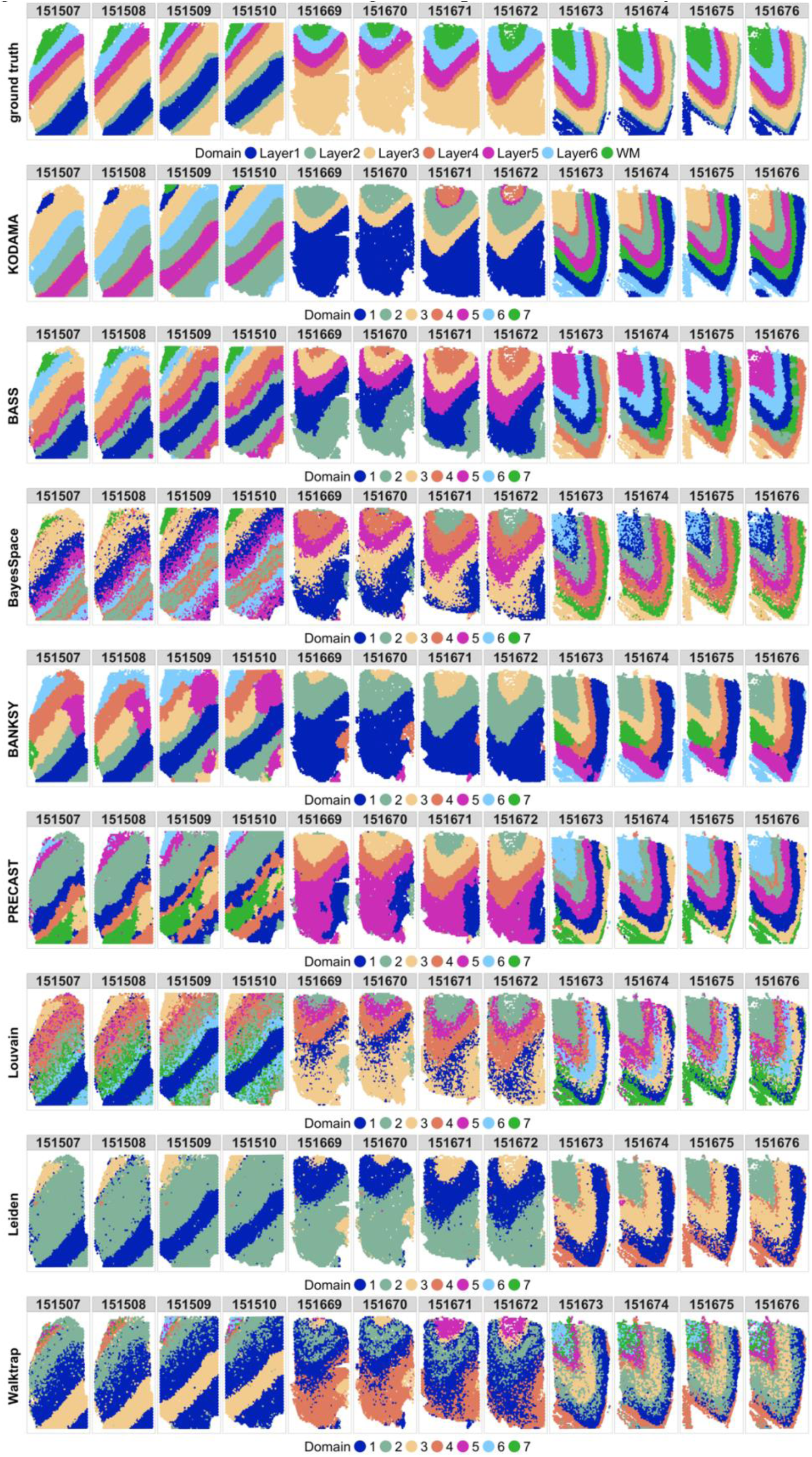
Clustering results across DLPFC tissue sections. Visualization of spatial clustering outputs for different algorithms on DLPFC tissue, emphasizing the performance of KODAMA in recapitulating known cortical layers.

**Supplementary Figure 9:**
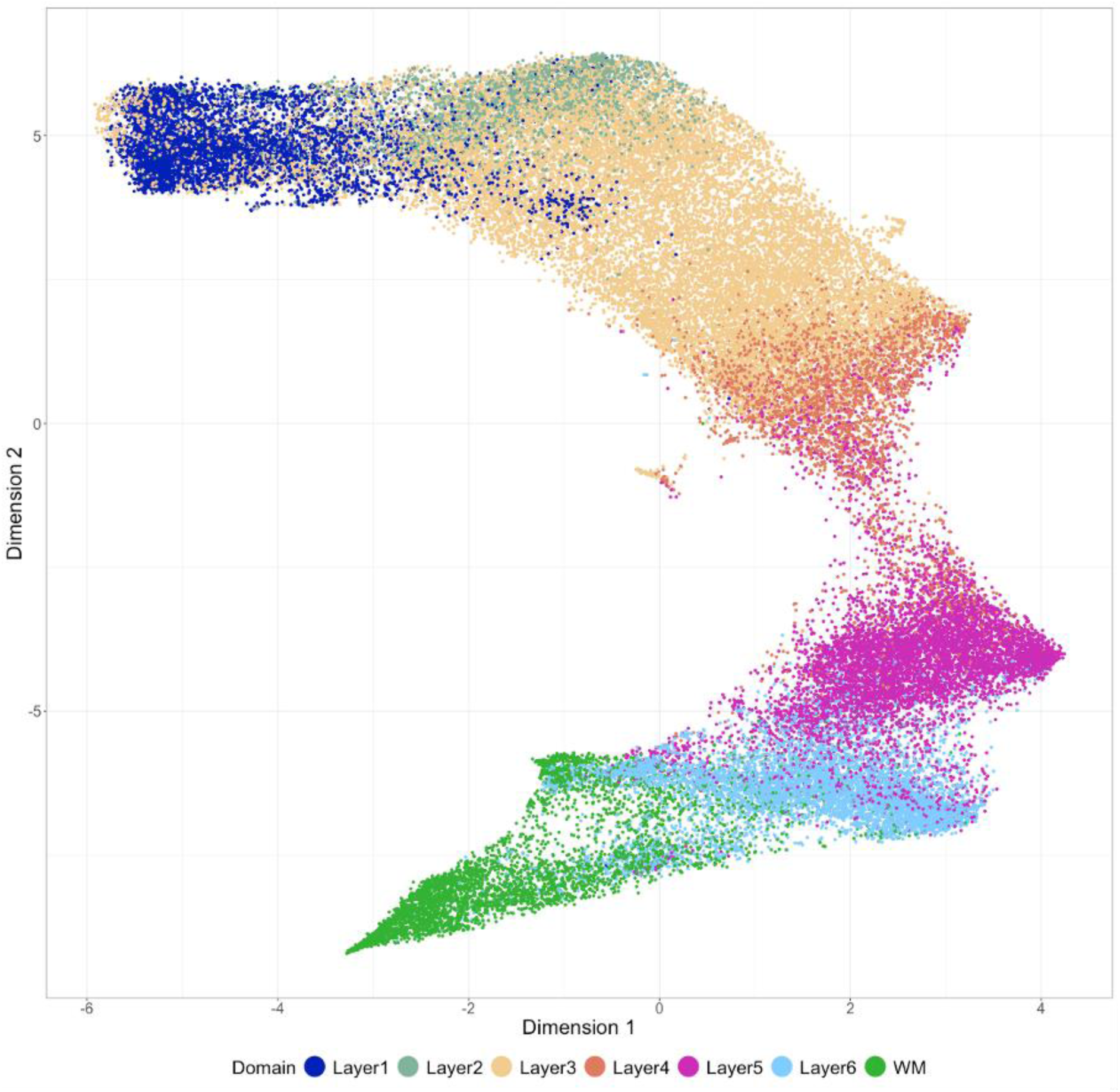
KODAMA analysis across all 12 DLPFC slides. Integrated clustering output using KODAMA across all sections, revealing fine-grained sublayer structures and spatial coherence between adjacent slices.

**Supplementary Figure 10:**
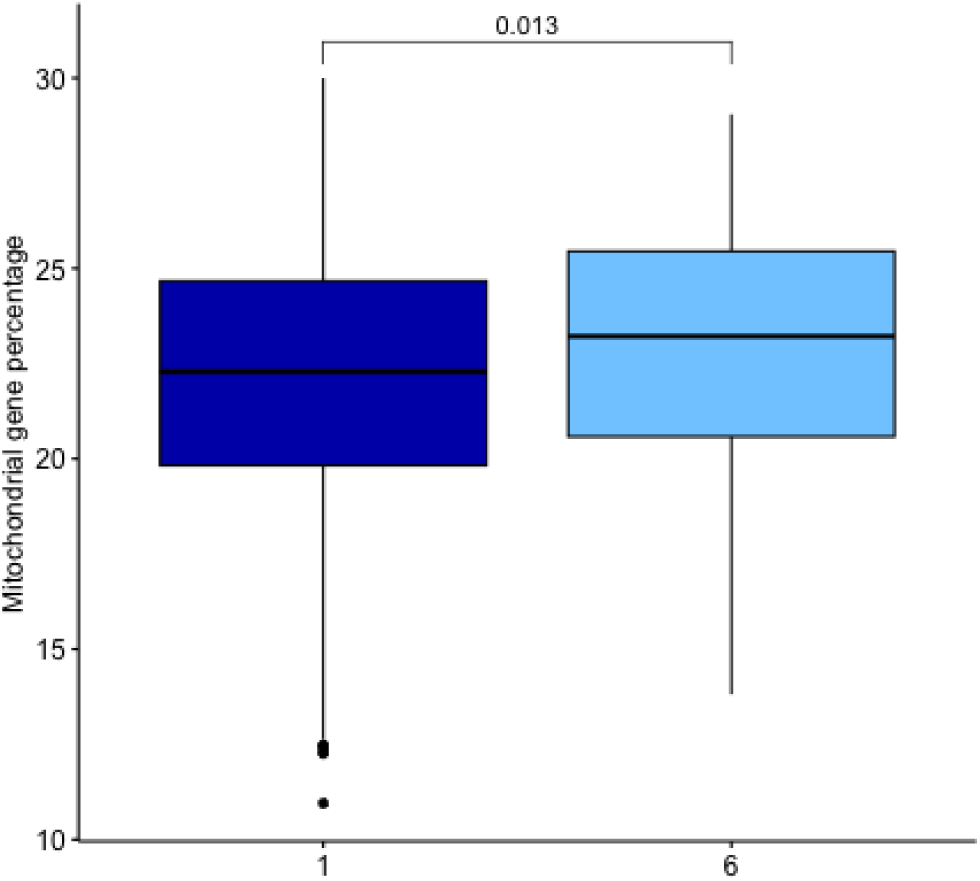
Mitochondrial content in outlier clusters. Comparison of mitochondrial gene expression in identified outlier clusters from DLPFC slides, highlighting regions with high mitochondrial content associated with tissue artifacts or stress zones.

**Supplementary Figure 11:**
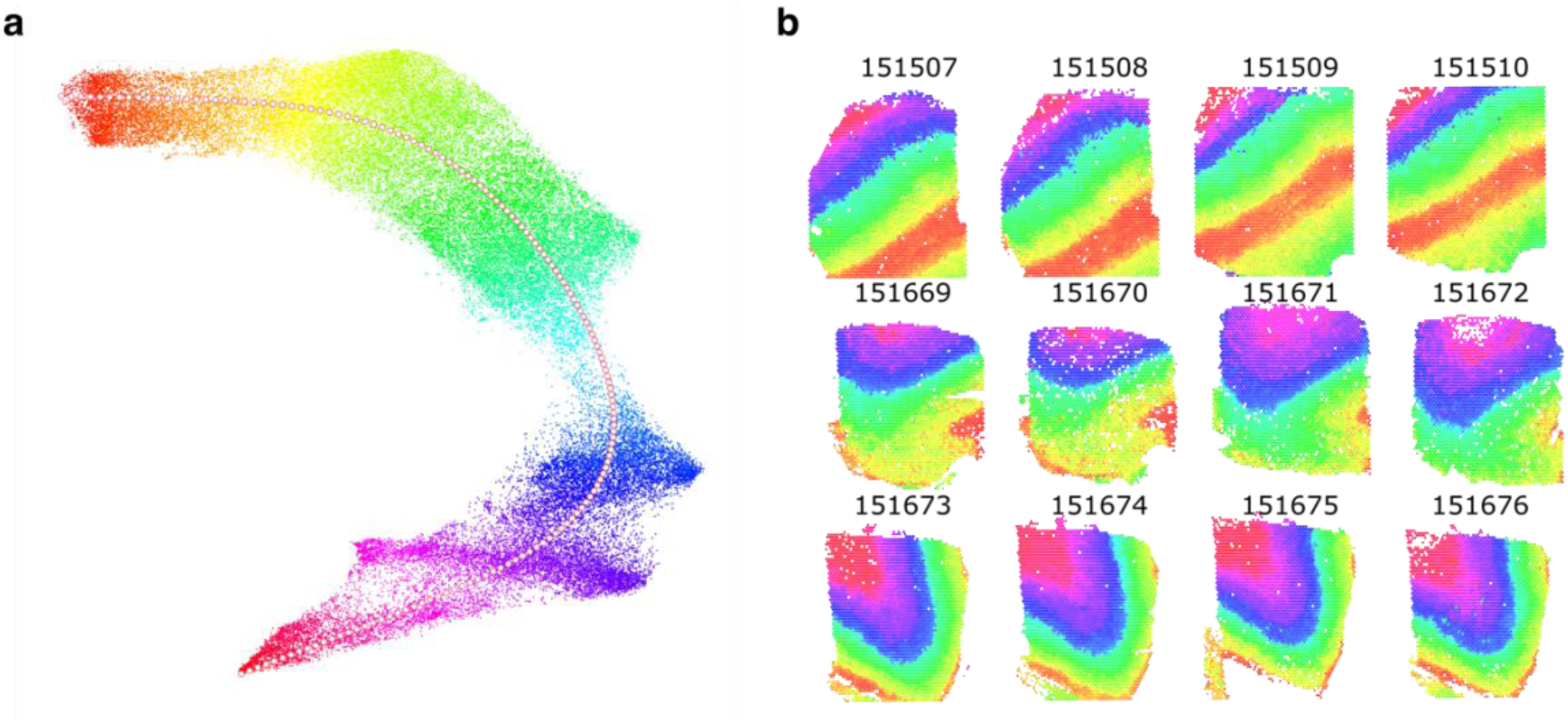
RGB trajectory in KODAMA dimensional space. RGB mapping of gene expression trajectories in **a,** KODAMA output, and **b,** slides of DLPFC dataset, enhancing the interpretation of spatial transitions.

**Supplementary Figure 12:**
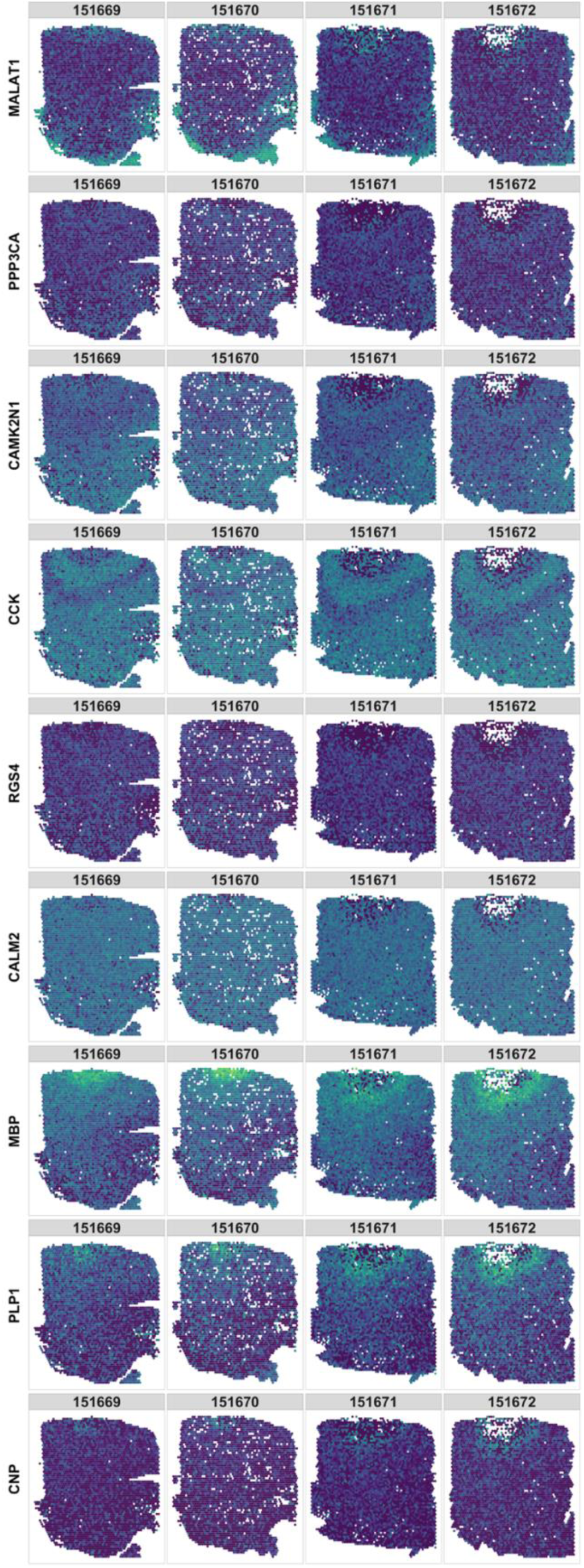
Gene projection validation. Further visualization of selected genes, confirming alignment with known anatomical features and demonstrating gradient continuity.

**Supplementary Figure 13:**
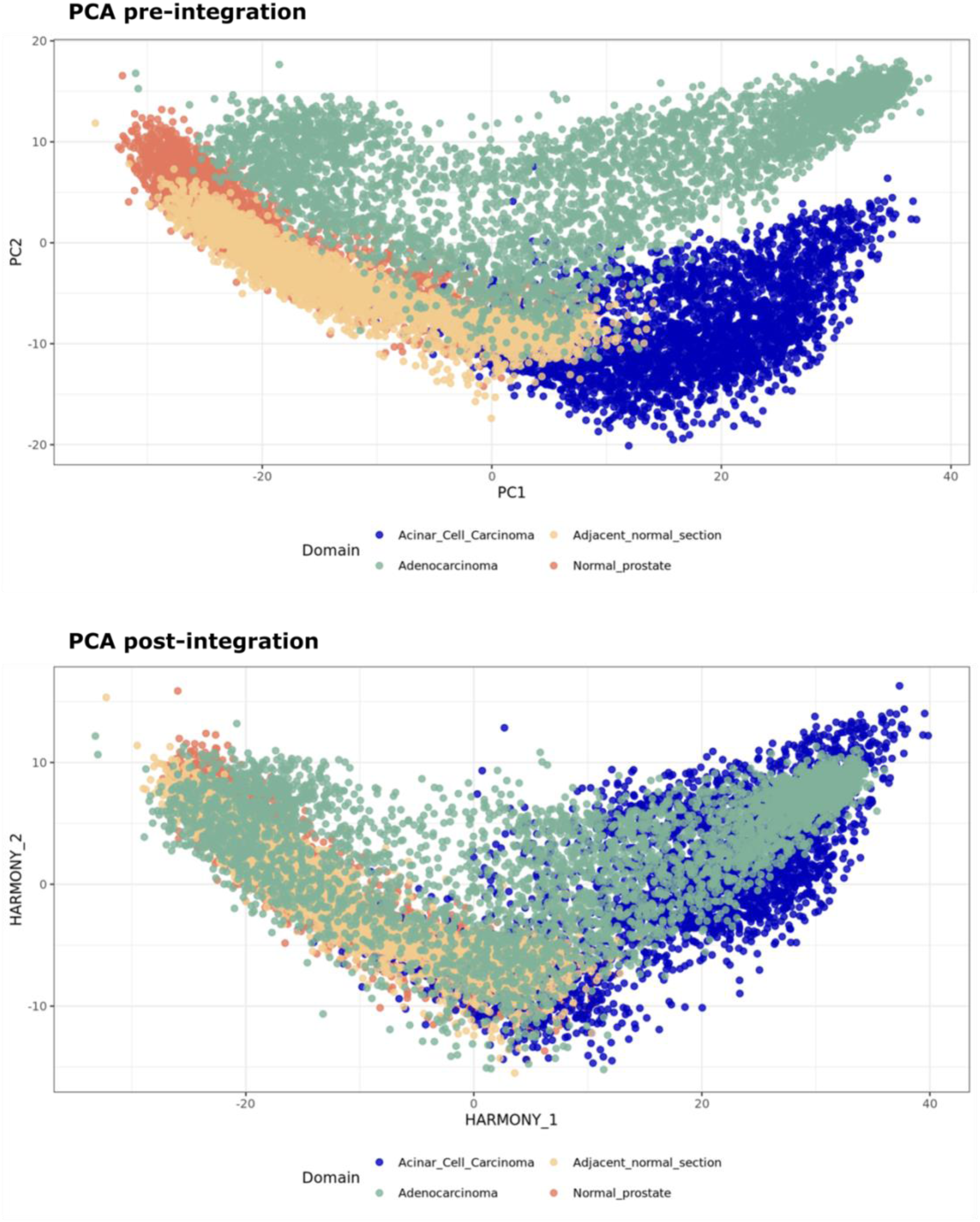
Pre- and post-integration visualization using PCA. Comparison of tissue variability across four prostate samples before and after Harmony integration using PCA, showing harmonized clustering and biological signal preservation.

**Supplementary Figure 14:**
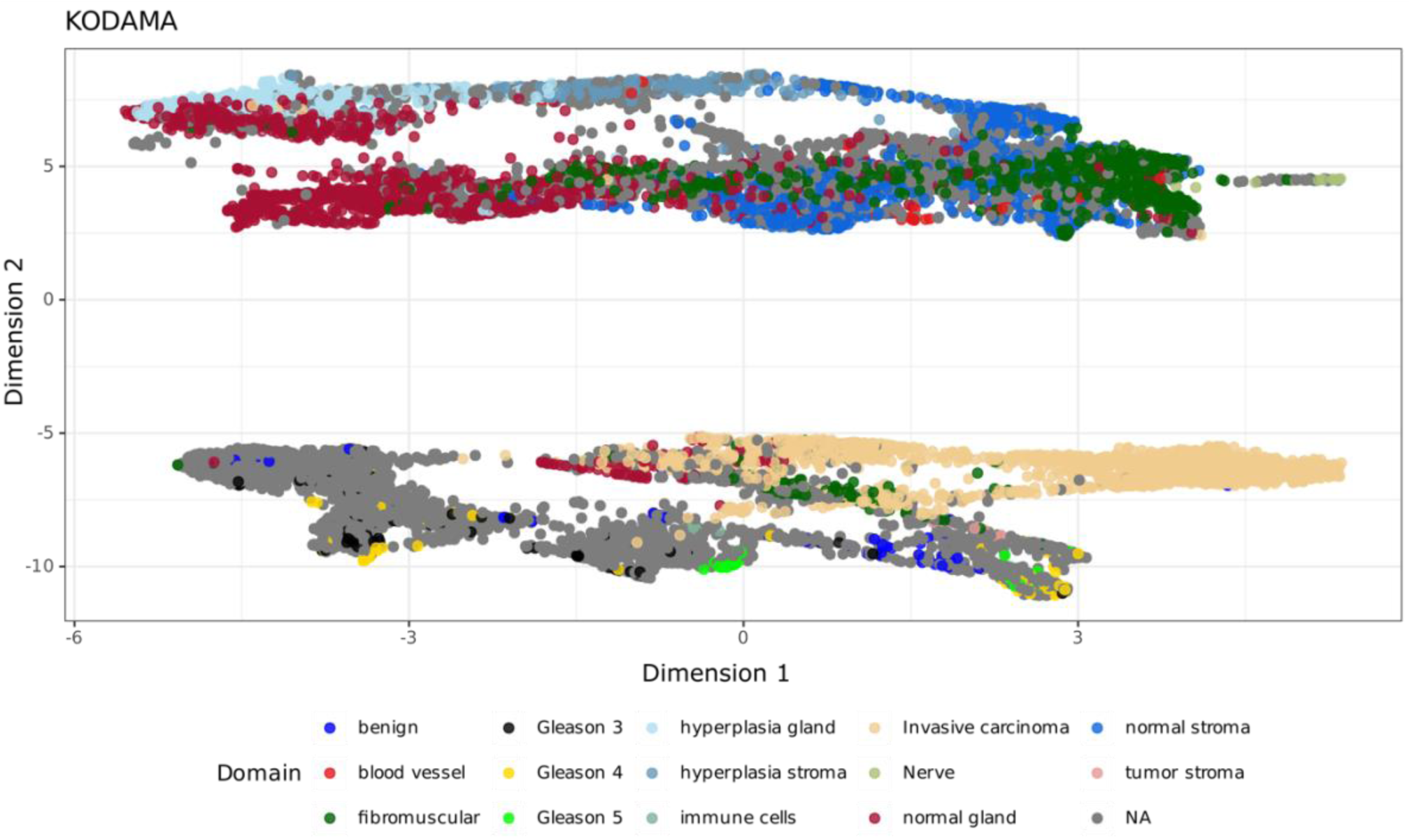
KODAMA analysis of prostate tissues. Visualization of pathology assignment on KODAMA output, identifying BPH zones and regions with tumor-like gene expression profiles.

**Supplementary Figure 15:**
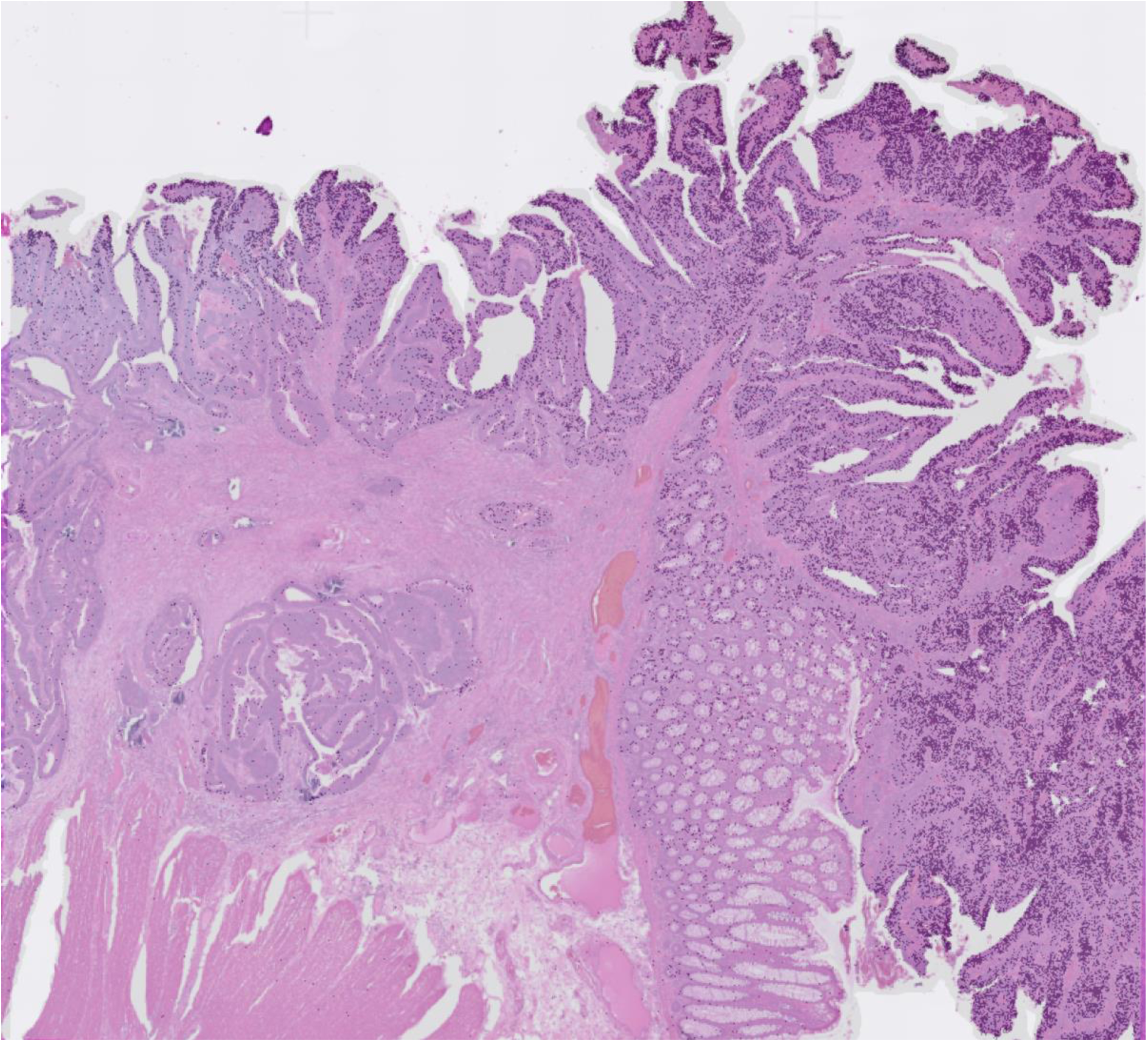
CXCL2 gene expression in colorectal tumor tissue.

**Supplementary Figure 16:**
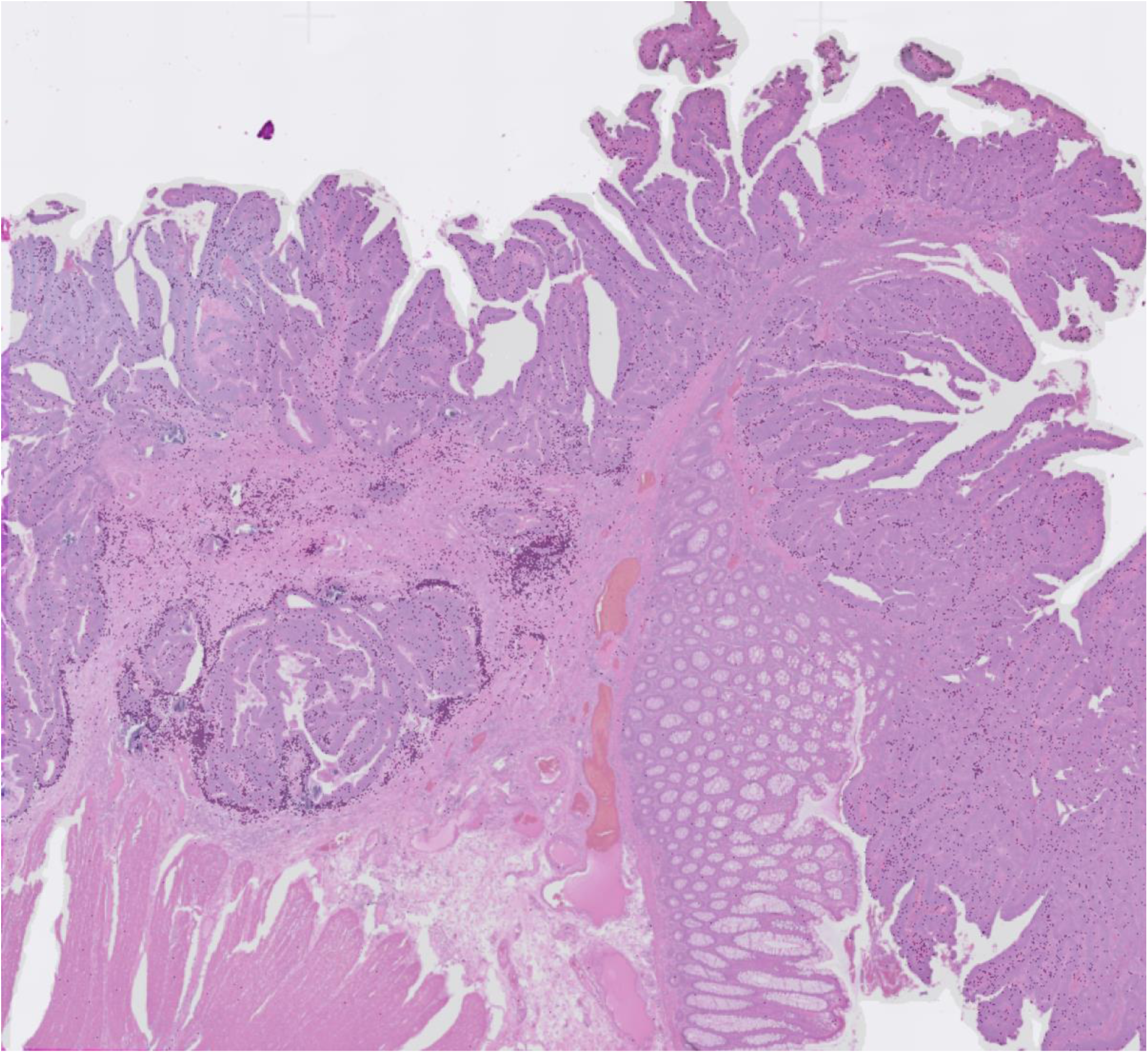
MMP11 gene expression in colorectal tumor tissue.

**Supplementary Figure 17:**
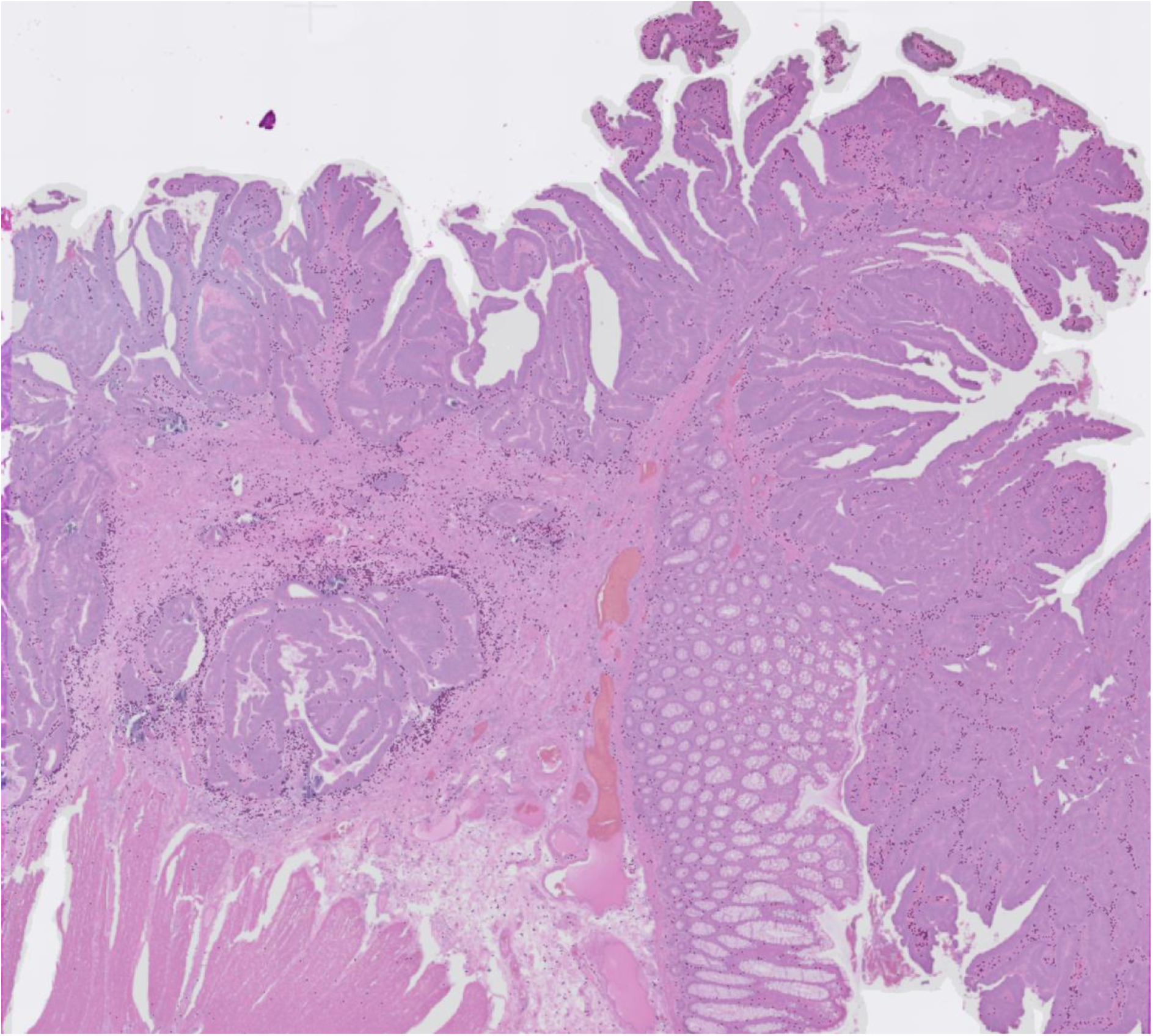
HTRA3 gene expression in colorectal tumor tissue.

**Supplementary Figure 18:**
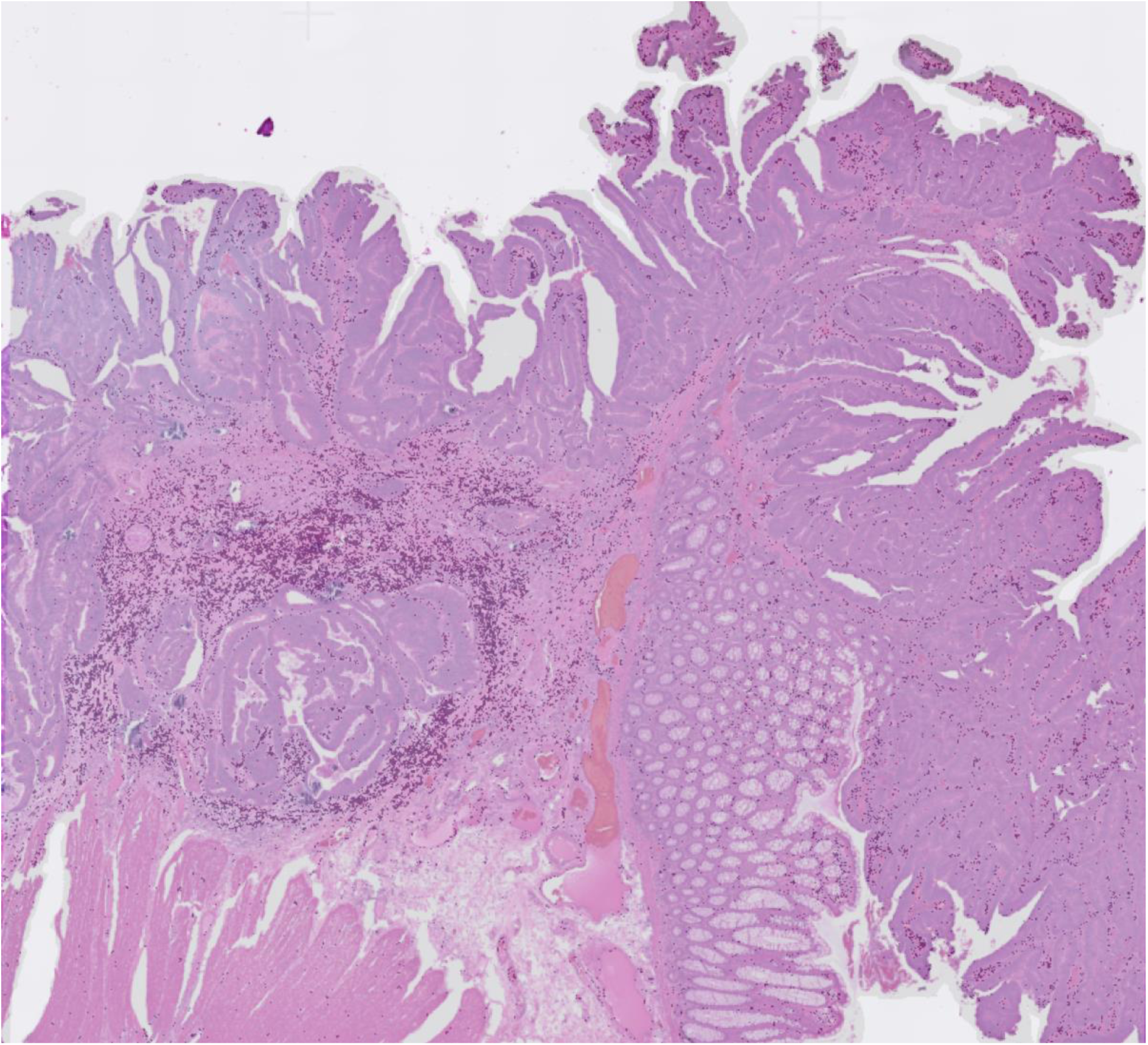
IGFBP3 gene expression in colorectal tumor tissue.

**Supplementary Figure 19:**
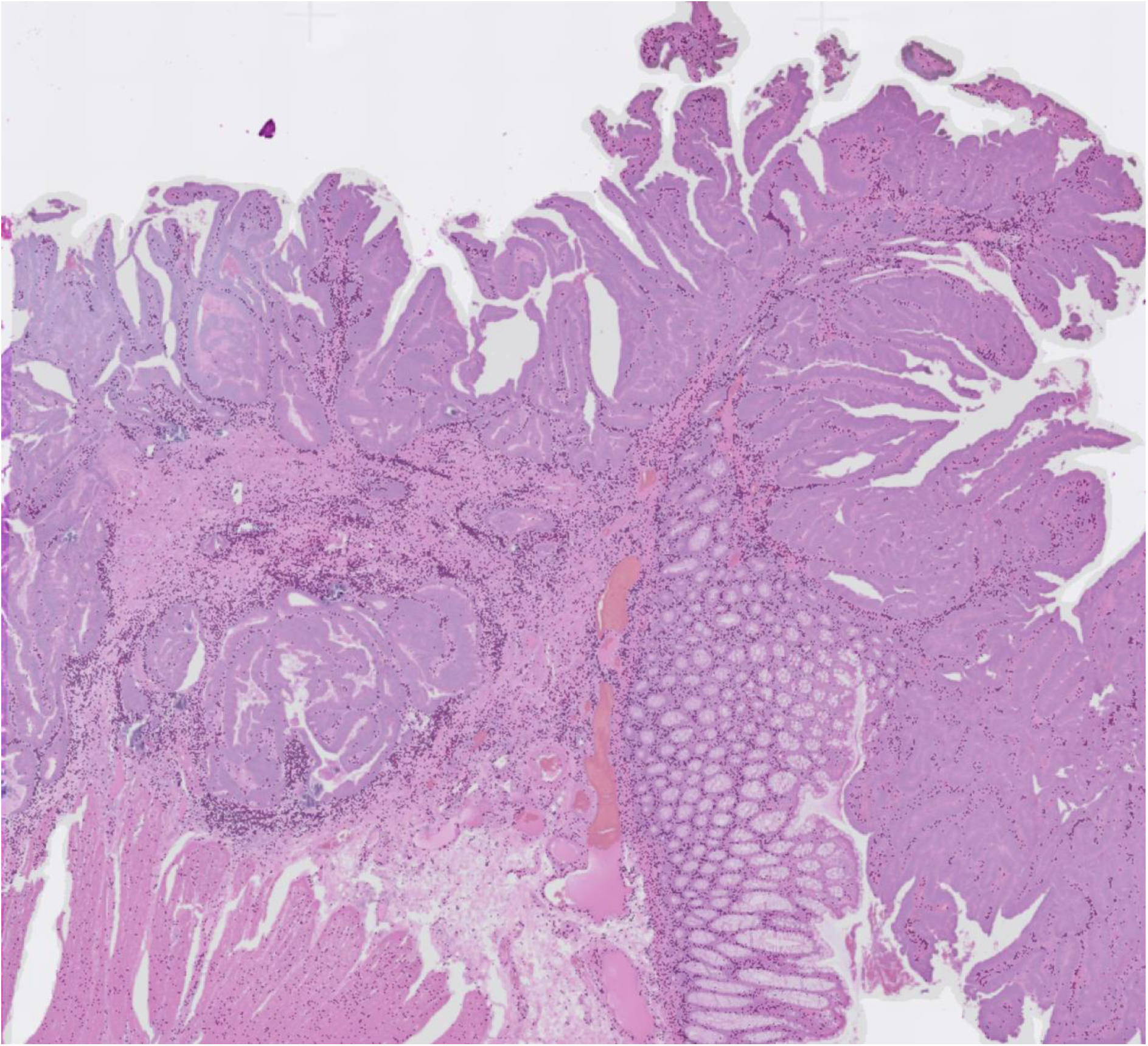
IGFBP5 gene expression in colorectal tumor tissue.

**Supplementary Figure 20:**
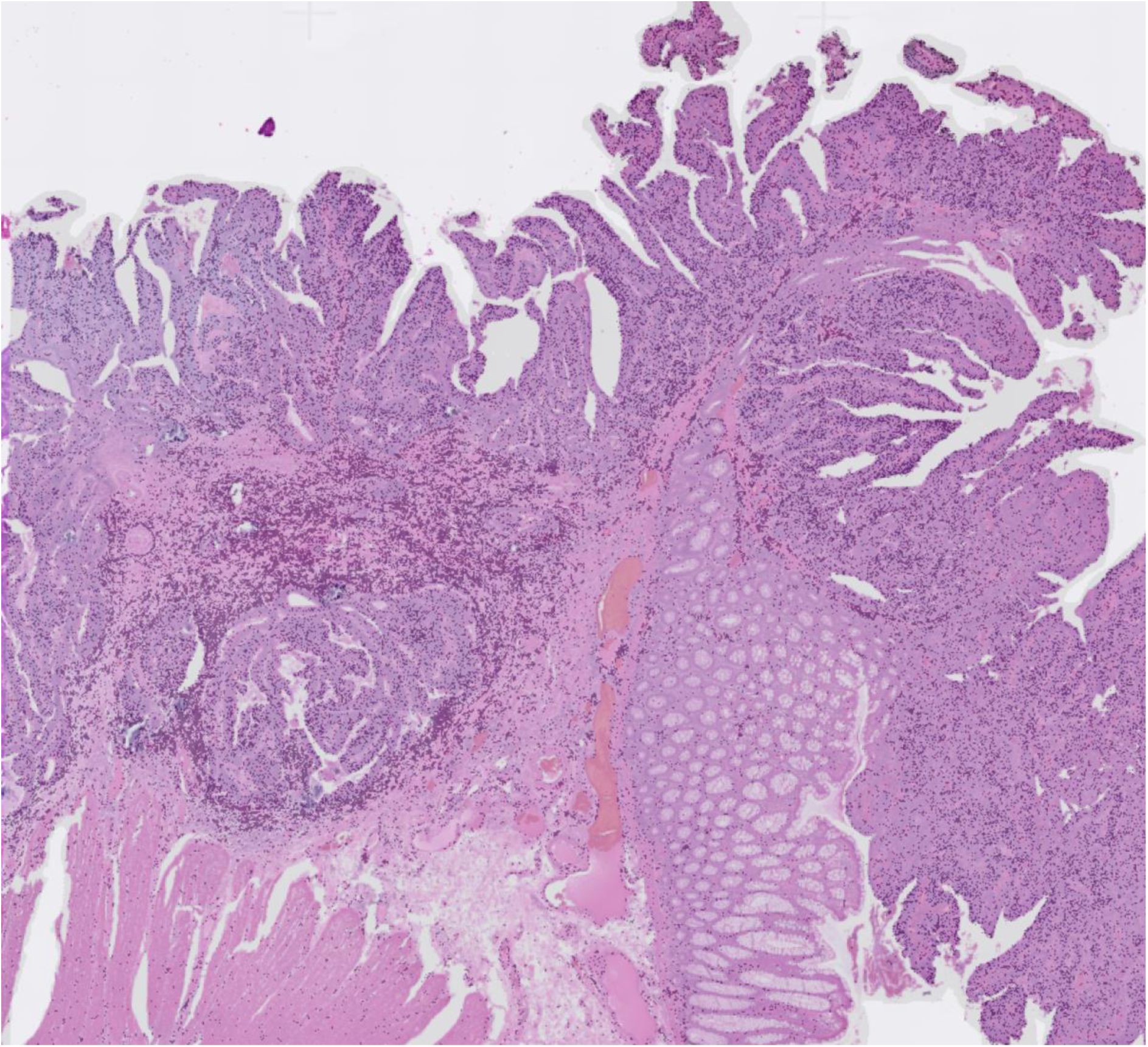
TIMP3 gene expression in colorectal tumor tissue.

